# Targeting the MAT2A-SAM axis increases platinum sensitivity in ovarian cancer by impairing DNA repair and stemness

**DOI:** 10.64898/2026.09.22.753550

**Authors:** Shu Zhang, Zhen Fu, Yanchi Zhou, Tara X. Metcalfe, Truc T. Vuong, Lindsay W. Brubaker, Benjamin G. Bitler, Heather M. O’Hagan, Kenneth P. Nephew

**Author notes:** Corresponding author: Kenneth P. Nephew, PhD, 302 Biology Building, Indiana University School of Medicine-Bloomington 1001 E. 3rd St, Bloomington, IN 47405.

## Abstract

Metabolic and epigenetic reprogramming drives platinum resistance and recurrence in high-grade serous ovarian cancer (HGSC), presenting a critical clinical challenge. Acquired platinum resistance has been linked to altered DNA methylation, a process requiring S-adenosylmethionine (SAM), synthesized by methionine adenosyltransferase 2A (MAT2A) from methionine metabolism. However, how the MAT2A-SAM axis coordinates metabolic-epigenetic remodeling in HGSC remains unclear. We demonstrate that MAT2A-driven SAM synthesis is essential for platinum-induced DNA methylation dynamics and ovarian cancer stem cells (OCSCs) enrichment. High MAT2A expression correlates with poor progression-free survival in HGSC patients, while genetic knockdown or pharmacological inhibition of MAT2A increases cisplatin sensitivity in HGSC cells. Genome-wide DNA methylation profiling showed that cisplatin reshaped the methylome with prominent hypermethylation at promoter regions, which was reversed by MAT2A knockdown. Transcription factors predicted from differential methylation were enriched in platinum resistance, DNA repair, and stemness pathways. Mechanistically, disrupting MAT2A-SAM axis suppresses a novel SAMTOR-mTOR-S6K-FANCD2 signaling pathway. This blockade leads to the accumulation of R-loops, attenuates both homologous recombination (HR) and non-homologous end joining (NHEJ) repair pathways, and enhanced platinum-induced cell death. Furthermore, transcriptomic analysis of paired primary and recurrent HGSC tumors links MAT2A increase to elevated OCSC features during recurrence. Consistently, MAT2A inhibition prevents cisplatin-increased OCSC population and sphere-forming capacity in vitro, both of which are rescued by exogenous SAM supplementation. Together, our findings identify the MAT2A-SAM axis as a pivotal metabolic-epigenetic regulator of the DNA damage response and cancer stemness, establishing MAT2A inhibition as a therapeutic strategy to enhance platinum sensitivity in HGSC.

**Highlights:**

- Cisplatin remodels the HGSC cell methylome with predominant promoter hypermethylation
- MAT2A-SAM is essential for platinum-induced methylome remodeling and OCSC enrichment
- MAT2A loss impairs activation of HR and NHEJ repair via mTOR-S6K-FANCD2 signaling
- MAT2A inhibition increases platinum sensitivity in both MTAP-proficient and -deficient HGSC cells tested

## Introduction

Platinum resistance remains a major therapeutic challenge in ovarian cancer and other cancers^1^. Most advanced-stage high grade serous ovarian cancers (HGSCs) respond well to upfront platinum-based chemotherapy^1^. Despite high initial response rates, the majority of HGSC tumors eventually acquire platinum resistance and recur. Recurrent HGSC is generally considered a fatal disease^2,3^. The development of resistance is a complex, multifactorial process involving genetic, epigenetic, metabolomic and other molecular programs that, despite intense investigation, remain poorly understood^4–6^.

HGSC tumors that respond poorly to chemotherapy have significant alterations in amino acid metabolism^7,8^, and metabolic profiling of platinum-resistant HGSC cell lines revealed increased methionine (Met) and cystine metabolism^9^. Met is an essential amino acid, and cancer cells exhibit an elevated demand for Met, a metabolic vulnerability known as methionine dependence or the Hoffman effect^10,11^. Moreover, recombinant methioninase (rMETase), which degrades Met, and low-Met intake effectively reduced tumor burden across multiple preclinical and clinical trials and sensitized tumors, including HGSC^12,13^, to both chemotherapy and radiation^14–16^. Central to Met metabolism, methionine adenosyltransferase 2A (MAT2A) converts Met to S-adenosylmethionine (SAM), the universal methyl donor used by a plethora of methyltransferase enzymes^11^. MAT2A inhibitors show selective synthetic lethality in methylthioadenosine phosphorylase (MTAP)-deficient cancers by preventing Met salvage and increasing dependence on de novo SAM synthesis^17–19^. In addition, byproducts of SAM-dependent transmethylation flux into the transsulfuration pathway, generating reduced glutathione (GSH), which facilitates redox balance and drug detoxification^20,21^, suggesting that MAT2A-driven SAM synthesis could be a therapeutic target to improve chemosensitivity.

A key phenomenon underlying disease relapse and recurrence in most cancers is the persistence of cancer stem cells (CSC)^22,23^. Although chemotherapy is effective at initially decreasing the tumor bulk, treatment leaves behind residual ovarian CSCs (OCSCs) capable of regenerating tumors and driving recurrence^24^. Understanding the mechanism(s) responsible for OCSC survival and developing strategies to target these residual cells are critical to blocking acquired resistance to chemotherapy and, thus, OC recurrence. We and others have shown that platinum-based chemotherapy induces epigenetic and metabolic reprogramming, enriching ovarian cancer stem cells (OCSC)^22,25,26^. Aberrant DNA hypermethylation of CpG islands in key genes regulates stemness and platinum resistance^27,28^, and DNA methyltransferase inhibitors (DNMTis) block platinum-induced OCSC enrichment to overcome platinum resistance, demonstrating that DNA methylation plays a critical role in regulating platinum-induced cellular reprogramming ^29,30^. In breast and lung cancer CSCs, Met and SAM availability has been shown to be a vulnerability by regulating histone methylation^31,32^. However, a unifying mechanism to explain how MAT2A-driven SAM synthesis coordinates platinum-induced DNA methylation remodeling and OCSC enrichment has not been investigated in HGSC.

In this study, we identify MAT2A-driven SAM synthesis as a key metabolic-epigenetic regulator in HGSC. Inhibiting MAT2A reshapes the methylome, enhances platinum sensitivity, and prevents platinum-induced enrichment of OCSCs. Methylome profiling analysis further demonstrates the critical importance of SAM synthesis in platinum-induced methylome alterations in HGSC. Building on this finding, we demonstrate that MAT2A-driven SAM synthesis mediates activation of the DNA damage response in HGSC cells through the SAMTOR-mTOR-S6K-FANCD2 axis. Functionally, exogenous SAM promotes platinum-induced OCSC enrichment, despite MAT2A inhibition, and notably, MAT2A inhibition is highly effective in both MTAP-deficient and -proficient HGSC cell lines. Together, our study reveals a previously unrecognized mechanism linking metabolism to epigenetic regulation through platinum-induced remodeling and, for the first time, demonstrates that platinum directly induces genome-wide DNA methylation changes in HGSC cells. Furthermore, our study establishes the MAT2A-SAM axis as a potential therapeutic target to enhance platinum sensitivity by abrogating DNA damage response and OCSC enrichment, ultimately to reducing OCSC-driven disease recurrence in HGSC.

## Methods

### Cell Lines and Cell Culture

HGSC cell lines were obtained from ATCC (OVCAR3, OVCAR5), and the Japanese Collection of Research Bioresources Cell Bank (Kuramochi). OVCAR3 and Kuramochi were maintained in RPMI 1640 supplemented with 10% FBS and 5% sodium pyruvate. OVCAR5 was maintained in DMEM (Gibco. cat.11965092) supplemented with 10% FBS. All cell lines were tested for mycoplasma contamination (Lonza, cat. #LT07-318) every 6 months and used within passages 2-15 after thawing. For methionine-depleted culture conditions, methionine-free RPMI 1640 (Gibco, cat. A1451701) was obtained from Thermo Fisher. Methionine-free DMEM was prepared from a base formulation lacking glutamine, methionine, and cystine (Gibco, cat. 21013024) by supplementing with L-glutamine (584.0 mg/L) and L-cystine (63.0 mg/L) as indicated in medium protocol. Information for plasmids, chemical compounds and drugs can be found in Supplemental Table S1.

### Tumor Cell Growth and Spheroid Formation Assay

IC_50_ of FIDAS-5 was determined and calculated as previously described^33^. HGSC cells were treated with ½ IC_50_ of platinum (cisplatin) for 3 hours (OVCAR3, 7.5 μM; OVCAR5, 6 μM;) and/or IC_50_ of FIDAS-5 (OVCAR3 11.8 μM; OVCAR5 10.9 μM). For MAT2A knockdown experiments, siMAT2A-treated cells were incubated for 24 hours, siRNA was removed, and cells were then treated with platinum for 3 hours. Following treatments, cells were seeded for growth assays (2,000 cells/well, 96-well plates), assessed daily using MTT (2 mg/mL) over five days and analyzed in GraphPad Prism. For spheroid formation, 2,000 (OVCAR3) or 1,500 (OVCAR5) cells/well were cultured in stem cell medium (24-well low attachment plate), and spheroid area was quantified after 10 days using ImageJ.

### Liquid Chromatography-Mass Spectrometry (LC-MS)

Cells were treated with IC_50_ platinum for 3hr, 8hr, or 16hr, washed with ice cold PBS, and flash-frozen at −80 °C. Intracellular Met, SAM, and S-adenosylhomocysteine (SAH) were quantified by targeted mass spectrometry performed by the Van Andel Institute Mass Spectrometry Core. Metabolites were extracted using a 4:4:2 (v/v/v) acetonitrile:methanol:water solution. Spike-in controls were prepared to confirm compound identification and serve as positive controls. Relative quantitation was performed on an Agilent 6470 triple-quadrupole mass spectrometer operated in ESI-positive dynamic multiple reaction monitoring (dMRM) mode, with chromatographic separation on a reversed-phase Waters T3 column. Data analysis was performed using Agilent MassHunter Quantitative Analysis software (v9.0).

### Xenograft and Immunohistochemistry (IHC)

Female NSG mice bearing subcutaneous OVCAR3 (5×10^6 cells) HGSC xenografts were randomized to receive vehicle or cisplatin (2 mg/kg, intraperitoneally, weekly, three weeks). This animal study was performed under Indiana University IACUC protocol #25037 and in accordance with guidelines approved by the Institutional Animal Care and Use Committee of Indiana University (Indianapolis, IN, USA). At the experimental endpoint, tumors were harvested, fixed in 10% neutral-buffered formalin, embedded in paraffin (FFPE), and sectioned by the Histology lab Service Core facility at Indiana University (Indianapolis, IN, USA). FFPE tissue sections were deparaffinized in xylene, rehydrated through graded ethanol, and subjected to heat-induced epitope retrieval in sodium citrate buffer (pH 6.0) for 3 minutes under pressure. After quenching endogenous peroxidase in 3% H_2_O_2_, sections were blocked with 5% goat serum and 1% BSA and incubated overnight at 4^◦^C with primary antibodies (MAT2A, ALDH1A1 and SIRT1). Signal was detected using SignalStain Boost Detection Reagent and DAB chromogen (CST, cat 11725). Slides were counterstained with hematoxylin, dehydrated, and mounted. Whole-slide images were captured using Motic EasyScan Pro scanner and analyzed for cell detection and DAB staining intensity using QuPath software.

### Bioinformatics Analysis

#### MAT2A analysis

MAT2A abundance was evaluated using CPTAC ovarian cancer proteomics data (cprosite)^34^. Log-transformed protein abundance were compared between tumor and normal ovarian tissue by Mann-Whitney U test (tumor n=84, normal n=19) for whole group and Wilcoxon signed-rank test (matched pairs n=10) for paired samples.

#### Single-Sample Gene Set Enrichment Analysi

*(ssGSEA) of paired primary/recurrent HGSC tumors*: ssGSEA was performed on 24 matched primary/recurrent pairs HGSC tumors (GSE295041)^8^ using Python gseapy^35^ (v1.1.12) with weighting exponent α=0.25 and rank normalization enabled. MAT2A expression change was quantified as ΔMAT2A=log (recurrent) - log (primary). Stemness enrichment was scored against a custom 37-gene CSC panel [Supplemental Table S2] and 10 MSigDB Hallmark gene sets selected for relevance to MAT2A-driven CSC biology^36^. Per-patient Δ ssGSEA scores were correlated with ΔMAT2A by Spearman rank correlation^37^, and pathway enrichment analysis across all 17 gene sets and 24 patients was visualized as a heatmap ordered by ΔMAT2A.

### qRT-PCR and Quantitative Methylation-specific PCR (qMSP)

Total RNA was extracted using the RNeasy Mini Kit (Qiagen, cat. 74104) and DNA was extracted using the DNeasy Blood & Tissue Kit (Qiagen, cat. 69504), each according to the manufacturer’s instructions. Nucleic acid concentrations were measured using a NanoDrop spectrophotometer. For gene expression analysis, 2 μg of total RNA was reverse transcribed using MAXIMA reverse transcriptase (ThermoFisher, cat. 3276916). For methylation analysis, 500 ng of DNA was bisulfite-converted using the EZ DNA Methylation Kit (Zymo Research, cat. D5031) following the manufacturer’s instructions. Methylation-specific primers were designed with MethPrimer. qRT-PCR and qMSP was performed with SYBR Green Master Mix (Roche, cat. 04887352001) on a Roche LightCycler 480. Relative gene expression was calculated as previously described^38^, with *EEF1A* or *RhoA* as the internal reference. Relative methylation levels were normalized to *Alu* and calculated using the comparative Ct method. Primers can be found in Supplemental Table S1.

### Enzyme-linked immunosorbent assay (ELISA)

Global DNA methylation was quantified using the MethylFlash™ Global DNA Methylation (5-mC) ELISA Easy Kit (Colorimetric; Cat. No. P-1030; EpigenTek, Farmingdale, NY, USA). Briefly, 200 ng of purified genomic DNA was added to each well, and 5-methylcytosine levels were detected according to the manufacturer’s instructions with detection antibodies. Absorbance was measured at 450 nm using a microplate reader, and the percentage of global DNA methylation (%5-mC) was calculated from a standard curve according to the manufacturer’s protocol.

### Methylome profiling

Genome-wide DNA methylation profiling was conducted using the Infinium HumanMethylationEPIC v2.0 BeadChip platform (Illumina, California, USA), enabling precise measurement of methylation of over 935,000 CpG sites^39^. Raw data were processed using the SeSAMe R package (v1.24.0)^40^, which normalized signal intensities, corrected dye bias, and filtered low-quality probes. β-values (0=unmethylated, 1=fully methylated) were used to identify differentially methylated loci (DMLs) and differentially methylated regions (DMRs), applying a threshold of 0.1 and adjusted P-value < 0.05 with Benjamini-Hochberg correction^40^. CpG sites were annotated relative to CpG islands, shores (0-2 kb), shelves (2-4 kb), and open sea regions, and by proximity to gene features including TSS200, TSS1500, 5′UTR, first exon, and gene body^40^. Ingenuity Pathway Analysis (IPA) (QIAGEN Digital Insights, California, USA) was used to identify pathways of genes differentially methylated between groups.

To investigate biological pathways potentially affected by methylation changes at transcription factor regulatory sites, DMRs were analyzed by Probabilistic Genomic Concordance to identify enriched transcription factor binding sites (TFBS). Transcription factors with FDR <0.05 were mapped to validated target genes using the TRRUST v2 database and literature curation^41^. Pathway enrichment was assessed by one-sided Fisher’s exact test with Benjamini-Hochberg correction. Hypermethylated and hypomethylated TFBS were interpreted as predicting reduced and increased regulatory accessibility, respectively^42,43^.

### Apoptosis Assay

Apoptosis was evaluated using Annexin V/propidium iodide (PI) staining (Cat# 640914, Biolegend). Cells were collected by trypsinization, washed twice with cold PBS, resuspended in Annexin V binding buffer, and incubated with Annexin V-FITC and PI for 15 minutes at room temperature in the dark. Samples were analyzed using LSRII or CytoFLEX. Data were analyzed using FlowJo. Early apoptotic cells were defined as Annexin V□/PI□, late apoptotic cells as Annexin V□/PI□, and viable cells as Annexin V□/PI□.

### Comet Assay and Quantification

DNA damage was assessed using the alkaline comet assay kit (Cat# 4250-50-K, R&D) according to the manufacturer’s instructions. Briefly, treated cells were harvested, washed with cold PBS, and resuspended at 1 × 10□cells/mL. 5-10 μl cells were mixed with low-melting-point agarose and immediately spread onto comet assay slides. Slides were allowed to solidify at 4°C for 10-15 minutes in the dark. Solidified slides were then immersed in the provided lysis solution at 4°C for 1 hour or overnight, followed by incubation in freshly prepared alkaline unwinding solution (pH>13) for 20-30 minutes at room temperature. Electrophoresis was performed under alkaline conditions at 25 V for 20-30 minutes. After electrophoresis, slides were rinsed and dried under at 37°C. Then cells were stained with SYBR Green for 30 minutes at room temperature. Comets images were captured using NiE (Nikon) microscope. At least 50 cells per sample were analyzed using CometScore 2.0 software (TriTek Corp., Sumerduck). DNA damage was determined by measuring and quantifying tail moments.

### Plasmid Transfection

Cells were seeded and transfected at approximately 80% confluency. Plasmids (shMAT2A, Flag-DEPDC5, and pLCN DSB Repair Reporter (DRR)) were transfected using Lipofectamine 3000 (Cat# L300015, Invitrogen) following the manufacturer’s instructions. Briefly, plasmid DNA was diluted in Opti-MEM and mixed with lipofectamine reagent. After incubation for 10-15 minutes at room temperature, the mix was added to the cells and incubated for 24-48 hours before selection with appropriate drugs. For DRR plasmid, cells were treated with neomycin for 10 days. For knockdown experiments using shMAT2A, cells were treated with puromycin 3 days post-transfection and then analyzed by western blot. For Depdc5 expression, cells were collected 48-72 hours post-transfection and then assessed by western blot.

### DNA Damage Reporter System

The DNA damage reporter (DDR) assay was performed as described^44^. Briefly, HGSC DDR reporter cell lines were generated and selected with neomycin for 10 days. Cells were co-transfected with pCBASceI and pCAGGS DRR mCherry Donor EF1α BFP plasmids using Lipofectamine 3000, with optimized plasmid ratios, for 8-16 hours. Cells were cultured for an additional 24 hours, then trypsinized and washed by PBS. DNA repair pathway fluoresces signal across different cell lines were quantified by flow cytometry using CytoFLEX.

### R-loop Detection and Immunofluorescence

R-loops were detected by immunofluorescence using the S9.6 monoclonal antibody. Briefly, cells were seeded on glass coverslips, treated as indicated, washed with PBS, and fixed in ice-cold methanol (−20 °C) for 7 minutes, followed by rehydration in PBS. Cells were quenched with 0.1 M glycine and permeabilized with 0.2% Triton X-100. Coverslips were incubated with mouse anti-S9.6 and rabbit anti-γH2Ax antibody (1:100-1:250) for 1h at room temperature (OVCAR5) or overnight at 4°C (Kuramochi), followed by incubation with Alexa Fluor-conjugated secondary antibodies (1:1000) for 1 hour at room temperature in the dark. Nuclei were counterstained with DAPI, and coverslips were mounted using DAPI-containing mounting medium. Images were acquired using a 60× oil immersion objective on an SP8 confocal microscope (Leica) and only signal within the nucleus was analyzed using ImageJ/Fiji.

### Co-immunoprecipitation (Co-IP) and Western Blot

For Co-IP experiments, FLAG-DEPDC5 expressing cells were washed with PBS and collected by scraping. Cells were lysed in Co-IP lysis buffer on ice for 30 minutes, followed by centrifugation to collect the supernatant. The clarified lysates were incubated overnight at 4°C with pre-washed anti-FLAG magnetic beads under rotation. Beads were washed three times (15 min each, rotating), and bound proteins were eluted in 1.5× Laemmli buffer at 95°C for 10 minutes before separation by SDS-PAGE and western blot analysis. For whole-cell western blot, cells were lysed in 4% SDS after complete removal of PBS. Lysates were homogenized using QIAshredder columns, and total protein concentration was determined by BCA assay as previously described^38^. Equal amounts of protein from each treatment group were loaded onto Bio-Rad SDS-PAGE gels and analyzed by western blot. Antibodies can be found in Supplemental Table S1.

### ALDEFLUOR Assay and FACS Sorting

ALDH enzymatic activity was measured using the ALDEFLUOR assay kit (Cat#01700, Stemcell Technologies) per the manufacturer’s instructions^38,45^, with DEAB as a negative control for gating. ALDH+ populations were analyzed on LSRII (BD Biosciences) and CytoFlex (Beckman Coulter). ALDH+ and ALDH− cells were sorted by FACS using the SH800 (Sony).

### Statistical Analysis and Data Availability

All data are presented as individual data points with mean ± SD from three independent biological replicates unless otherwise indicated. Statistical comparisons between different treatment were performed using Student’s t-test or one-way ANOVA as needed. EPIC Array V2 data (GSE335264) are available at Gene Expression Omnibus data repository at the National Center for Biotechnology Information (NCBI).

## Results

### Methionine dependency and platinum-associated methionine-cycle remodeling in HGSC

Given the potential importance of targeting methionine (Met) metabolism in cancers, we first examined whether HGSC cells depend on Met by culturing HGSC cell lines in Met-depleted media and assessing cell growth. Over a period of five days, proliferation of HGSC cell lines [Fig.1A, Supplemental Fig.S1A and S1B] was inhibited by Met-depleted [Met(–)] compared to complete medium. Next, using an untargeted metabolomic dataset (ST002010)^46^ of platinum-resistant and -sensitive CaOV3 HGSC cell lines, we examined whether Met-related metabolites were specifically altered in association with platinum resistance. Levels of Met and SAM were unchanged between platinum-resistant and platinum-sensitive CaOV3 cells [Supplemental Fig.S1C]; in contrast, platinum-resistant cells exhibited increased levels of 5′-methylthioadenosine (5′-MTA), cystathionine, and glutathione (GSH), indicative of enhanced polyamine synthesis and transsulfuration, while decreased creatine and glycerophosphocholine (GPC) levels [Supplemental Fig.S1C]. These result suggested rapid SAM turnover in platinum-resistant cells and a selective redirection and maintenance of metabolic flux toward polyamine synthesis to support DNA stabilization^47^ and the transsulfuration pathway to produce GSH^21^.

**Figure 1.**
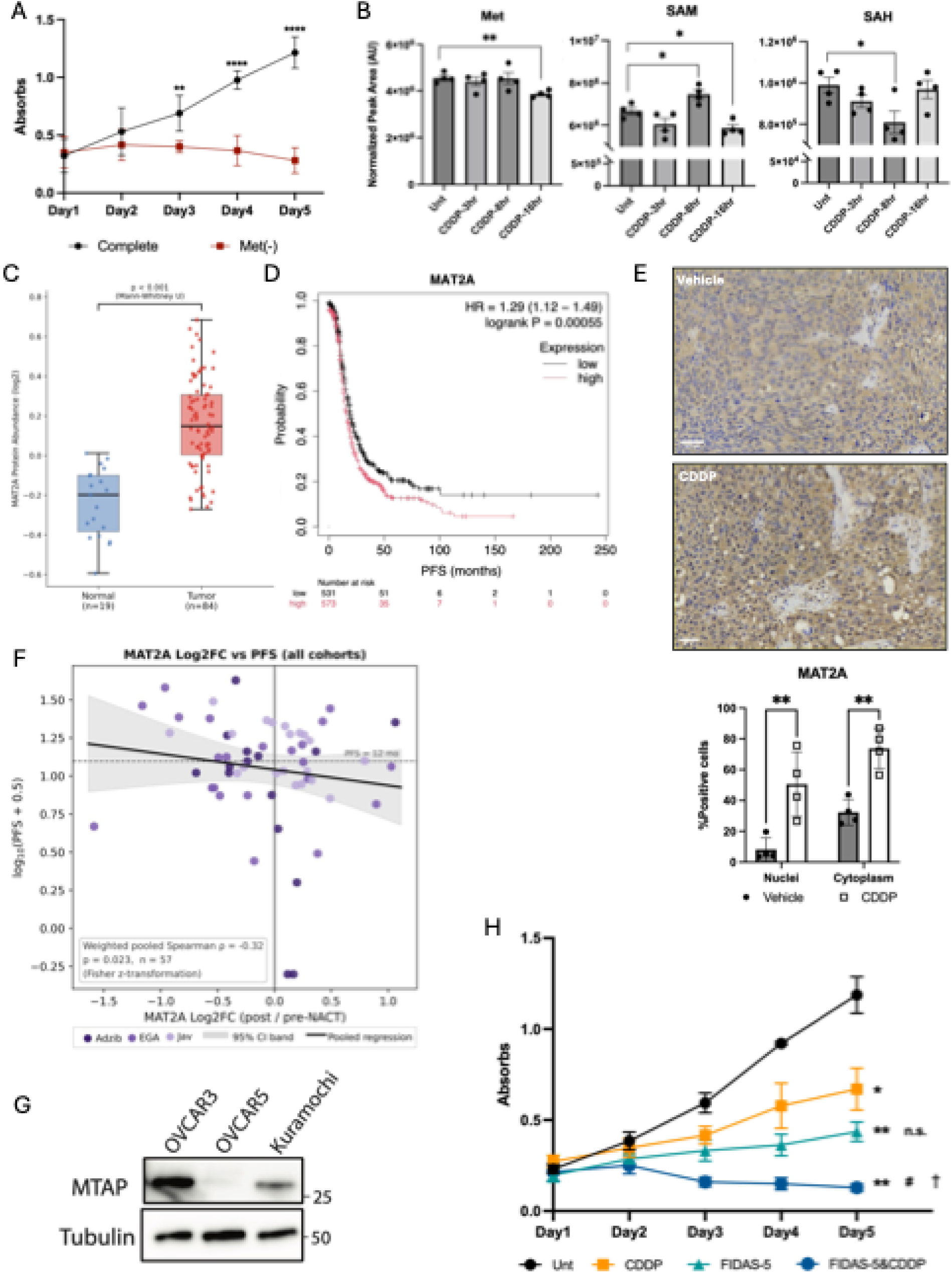
High MAT2A associates with poor prognosis, and MAT2A inhibition suppresses OC tumor growth. **(A)** Proliferation of OVCAR3 in complete or methionine-depleted [Met(–)] medium over five days. Data are presented as the mean ± SD of three biological replicates. **(B)** Intracellular levels of Met, SAM, and SAH were quantified by LC-MS and expressed as normalized peak area (AU) with time course platinum (cisplatin, CDDP) treatment. **(C)** MAT2A protein abundance in normal ovarian tissue versus HGSC tumor tissue from the CPTAC dataset. **(D)** Kaplan-Meier progression-free survival (PFS)^48^ curves stratified by MAT2A expression (high vs. low) in HGSC patients. **(E)** Representative immunohistochemical staining of MAT2A in xenograft tumors from vehicle- and CDDP-treated mice. MAT2A staining intensity was quantified using QuPath and is shown below. Scale bar = 50μm. Data are presented as mean ± SD. **(F)** Pooled Spearman correlations between MAT2A log2 fold-change (post-vs. pre-neoadjuvant chemotherapy (NACT)) and log-transformed progression-free survival (PFS) across three independent HGSOC cohorts (n = 57 pooled). **(G)** Western blot of MTAP in HGSC cell lines OVCAR3, OVCAR5 and Kuramochi **(H)** Proliferation assay of OVCAR3 with CDDP (7.5μM, 3 hours) and/or FIDAS-5 (11.8μM, 48 hours) treatment over five days. Data are presented as the mean ± SD of three biological replicates. Symbols (**\***, **#**, †) indicate statistical significance (*p*<0.05) for the following comparisons: **\***, Unt vs. other treatments; **#**, CDDP vs. FIDAS-5 or combination; †, FIDAS-5 vs. combination. Higher symbol densities (e.g., **, ***) represent*p*<0.01 and *p*<0.001, respectively. (* p<0.05, ** p<0.01, *** p<0.001, **** p<0.0001; ns, not significant)

To investigate whether platinum acutely alters Met metabolism core metabolites, we treated OVCAR3 cells with platinum (cisplatin) IC_50_ and measured Met, SAM, and SAH levels at 0, 3, 8, and 16 hours using LC-MS. The level of Met remained largely unchanged following cisplatin; however, SAM levels increased by 8 hours, then tapered off by 16 hours, and inversely, the level of SAH decreased by 8 hours, then returned to baseline by 16 hours [Figure 1B]. Taken together, these dynamics suggest that acute platinum exposure transiently activates the Met cycle rather than causing passive metabolites accumulation, whereas resistant cells maintain a hypermetabolic state with selective downstream utilization of Met pathway constituents [Supplemental Fig.S1C]. These data highlight SAM synthesis as a target to prevent platinum-induced reprogramming of the Met cycle.

### Altered MAT2A-driven SAM synthesis is associated poor clinical outcome in HGSC

We next assessed the clinical importance of MAT2A. We analyzed CPTAC proteomics data^34^ and a curated HGSC transcriptomic survival cohort^48^ and evaluated the impact of expression levels of the SAM-synthesizing enzyme on patient outcomes. MAT2A protein levels were higher in tumor tissues than in normal ovarian tissues, including in paired patient samples [Fig. 1C; Supplementary Fig. S1D], while MAT2A mRNA levels were higher in HGSC than in normal fallopian tube epithelium^49^ [Supplementary Fig. S1E]. Furthermore, high MAT2A expression was associated with poor progression-free survival (PFS) in HGSC patients [Fig.1D]. To determine whether platinum regulates MAT2A expression *in vivo*, we analyzed xenograft tumors from vehicle- and cisplatin-treated mice [Supplemental Fig.S1F]. Compared with vehicle, cisplatin-treated tumors showed significantly increased nuclear and cytoplasmic MAT2A staining [Fig.1E].

To determine the potential clinical relevance of platinum-induced MAT2A upregulation, we examined the correlation between post-neoadjuvant chemotherapy (NACT) MAT2A expression and PFS across three independent HGSC cohorts^50^ (Adzib^51^, EGA (EGAD00001006456), Jav^5^, n=57 total). Interestingly, we found that NACT upregulation of MAT2A negatively correlated with PFS [Fig.1F, pooled Spearman ρ=-0.32, p=0.023; Fig.S1G], suggesting MAT2A expression as a potential biomarker for NACT response and outcome in patients with advanced-stage HGSC.

MAT2A inhibitors induce selective lethality in MTAP-deleted tumors, as the deletion reduces the regeneration of Met therefore increasing dependence on MAT2A-driven SAM synthesis^17,19^. Although MTAP-mutations are relatively rare in HGSC in general, occurring in only about 13% of cases^52,53^, it was of interest to examine this possibility. In agreement with published reports^54,55^, the level of MTAP in OVCAR3 was high, MTAP expression in Kuramochi was moderate and no expression of MTAP was observed in OVCAR5, a known MTAP-deleted cell line [Fig.1G]. We next determined the IC□□of the MAT2A inhibitor FIDAS-5 (OVCAR3 11.8 μM; OVCAR5 10.9 μM, 48 hours) [Supplemental Fig.S1H-I] and used this concentration in MTT assays to examine whether MAT2A enzymatic inhibition reduced cell proliferation in MTAP-proficient (OVCAR3) and -deficient (OVCAR5) cells. FIDAS-5 treatment significantly suppressed growth of both HGSC cell lines, and the combination of platinum with FIDAS-5 further reduced cell proliferation compared to either agent alone [Fig.1H; Supplemental Fig.S1J], demonstrating that inhibiting MAT2A has broad anti-proliferative effects in both MTAP status, either alone or in combination with platinum.

### Inhibiting MAT2A abrogates platinum-induced hypermethylation in HGSC

Platinum chemotherapies induce aberrant gene-specific promoter DNA methylation^5,27^ and platinum resistance is associated with methylome alterations^25,28,56^. However, whether platinum directly induces global DNA methylation remodeling remains unknown. We observed that 8-hour platinum treatment increased the methylation index (SAM/SAH), which indicates the capacity for methylation reactions in cells [Fig.2A], and global 5-mC levels also increased by 16 hours [Fig.2B], demonstrating a global increase in DNA methylation following platinum treatment. Because DNA methylation depends on SAM availability and DNMT activity^11^, we hypothesized that MAT2A and DNMTs levels are correlated in HGSC. To test this hypothesis, we carried out single-sample gene set enrichment analysis (ssGSEA) of 24 paired primary and recurrent HGSC tumors^8^. ΔMAT2A positively correlated with both ΔDNMT1 (Spearman r = +0.564, p = 0.0041) and ΔDNMT3A (Spearman r = +0.621, p = 0.0012) [Fig.2C], further leading us to hypothesize that inhibiting MAT2A would abrogate the effects of platinum treatment on the DNA methylome. Before performing genome-wide analyses to test this hypothesis, we validated the effect of MAT2A siRNA knockdown on MAT2A mRNA and protein [Supplementary Fig.S2A-B] and on locus-specific DNA methylation by using methylation-specific qPCR (qMSP) analysis. Two genes known to be regulated by DNA methylation, *BRCA1* and *ALDH1L1*^57,58^ were tested. OVCAR3 cells were treated with siScramble (control), cisplatin (IC_50_, 16hr) and/or siMAT2A (48hr). Platinum-induced promoter DNA methylation on *BRCA1* [Supplemental Figures S2C] and *ALDH1L1* [Supplemental Figures S2D], was inhibited by siMAT2A treatment.

**Figure 2.**
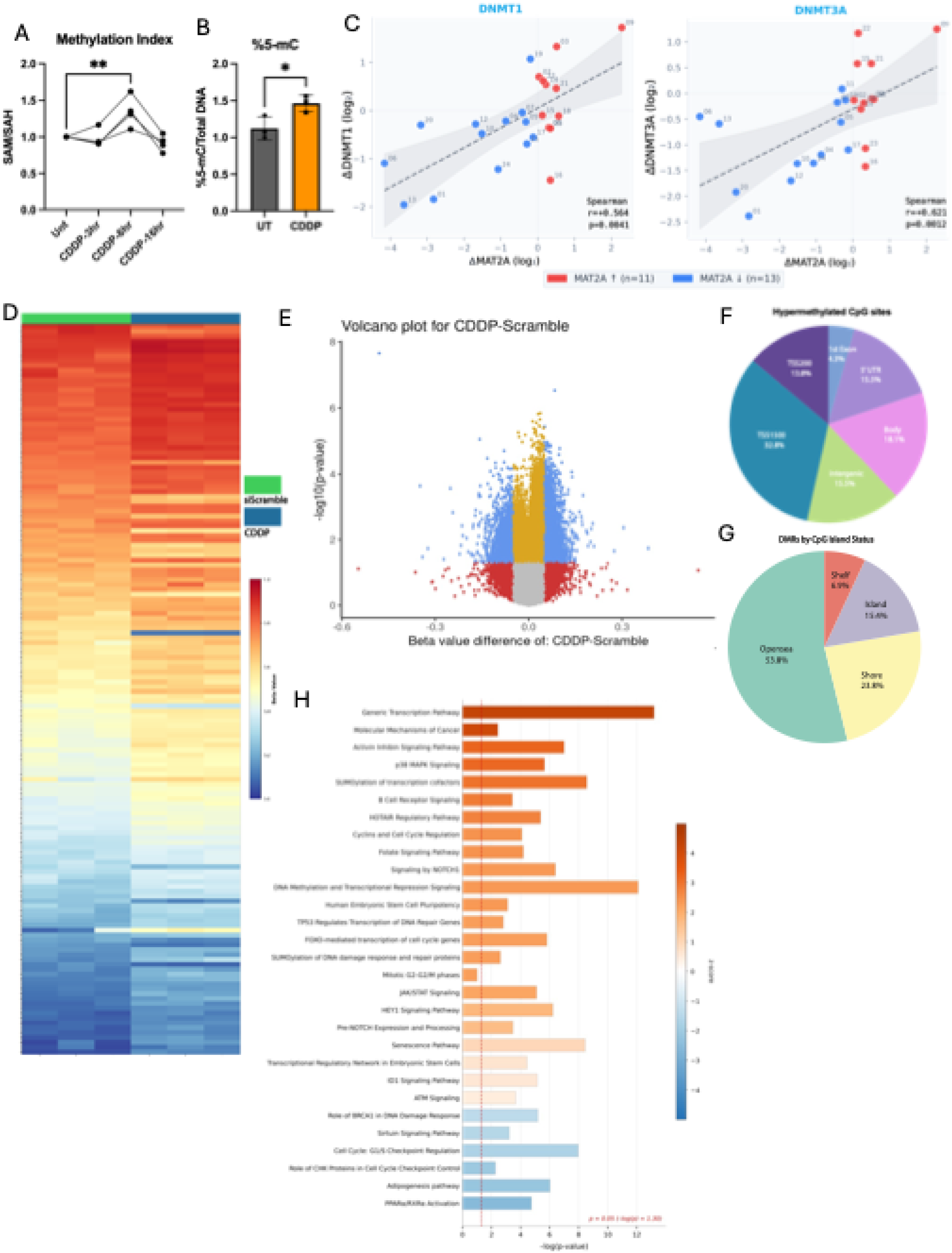
Platinum induces promoter-focused hypermethylation across the OC methylome. **(A)** Methylation index, calculated as the ratio of SAM to SAH), of OVCAR3 across a time course of platinum (cisplatin, CDDP) treatment (3, 8 and 16 hours) **(B)** Global 5-methylcytosine (5-mC) levels of OVCAR3 cells untreated (UT) or treated with CDDP (15μM, 16 hours) measured by ELISA and normalized to total level of DNA. **(C)** ssGSEA-derived ΔDNMT1 (left) and ΔDNMT3A (right) scores plotted against ΔMAT2A expression between matched recurrent and primary tumors stratified by MAT2A-high (red) and MAT2A-low (blue) patients. **(D)** Heatmap of beta values for statistically significant (adjusted p<0.05) differentially methylated probes identified between siScramble and CDDP-treated conditions. **(E)** Volcano plot of CpG-level methylation changes between CDDP-treated and siScramble control cells. Each point represents a single CpG site; blue points indicate statistically significant differential β-value (adjusted p < 0.05, Δβ> 0.05); red points indicate β-value changes below statistical significance; yellow points indicate statistically significant but Δβ-value < 0.05. Positive and negative β-value denote hypermethylation and hypomethylation in CDDP versus siScramble, respectively. **(F)** Pie chart of the distribution of hypermethylated CpG to gene features (promoter, exon, intron, and intergenic regions) in CDDP versus siScramble. **(G)** Pie chart of genomic annotation of differentially methylated regions (DMRs) across CpG island contextual categories (island, shore, shelf, and open sea) in CDDP versus siScramble. **(H)** Ingenuity Pathway Analysis (IPA) of biological pathways enriched among DMR-associated genes identified from the CDDP versus siScramble. (* p<0.05, ** p<0.01, *** p<0.001, **** p<0.0001; ns, not significant)

We next used the same treatment scheme as qMSP to investigate genome-wide changes at single-CpG resolution using the Infinium MethylationEPIC followed by analysis of the differentially methylated regions (DMRs). Principal component analysis (PCA) showed separations between the treatment groups [Supplementary Fig.S2G]. Hierarchical clustering of probes from platinum vs. siScramble revealed widespread gain of CpG methylation [Fig.2D] and CpG sites with positive Δβ-values [Fig.2E]. Hypermethylated CpG sites were markedly (approximately 46%) enriched by platinum at TSS200 and TSS1500 [Fig.2F], with 15.4% mapping to CpG islands [Fig.2G], suggesting “promoter-focused” DNA methylation changes. To assess concordance between platinum-induced methylomic and transcriptomic changes, DMRs were overlapped with differentially expressed genes (DEGs; RNA-seq data^30^), revealing that hypermethylated and downregulated group (Hyper+DOWN) comprised the largest proportion of DMR-DEG pairs [Supplemental Fig.S2H]. IPA of DMR-associated genes also identified enrichment of the “activation of DNA methylation and transcriptional repression signaling” pathway, supporting coordinated methylation-associated gene repression following platinum treatment [Fig.2H]. The enriched pathways also spanned several programs of direct relevance to platinum chemotherapy, including cell-cycle and checkpoint control (G1/S checkpoint regulation and cyclins), senescence pathways (p38 MAPK and oxidative-stress-induced senescence), and DNA damage/repair related pathways (TP53-regulated DNA repair genes, ATM signaling, and BRCA1 in DNA damage response) [Fig.2H, Supplementary Table S3]. Overall, these results are consistent with platinum altering DNA methylation in HGSC and provide insight into how these DNA methylome changes affect broad metabolic pathways as well as selective yet related cellular pathways in HGSC.

Having established that platinum induced genome-wide hypermethylation across the genome, we next investigated the effect of combining siMAT2A with platinum on DNA methylation, compared to platinum alone. A marked reduction in DNA methylation and a redistribution toward lower methylation level across CpG probes in hierarchical clustering [Fig.3A] and a negative shift of Δβ-values [Fig.3B, volcano plot] was observed in HGSC cells treated with siMAT2A + platinum combination compared to platinum only. Genomic location analysis further revealed that CpG sites hypomethylated in siMAT2A+platinum compared to platinum-only samples were enriched in TSS200-1500 (24.3%) and largely in intergenic regions (41.8%) and the majority of DMRs (86%) localized to open sea regions [Fig.3C-D], demonstrating that MAT2A inhibition resulted in a less promoter-focused methylomic landscape compared to platinum treatment. While these results support a role for MAT2A in the platinum-induced increase in DNA methylation, the overall number of pathways with significant activation or inhibition (|z| > 2) identified by IPA was limited [Supplemental Table S4].

**Figure 3.**
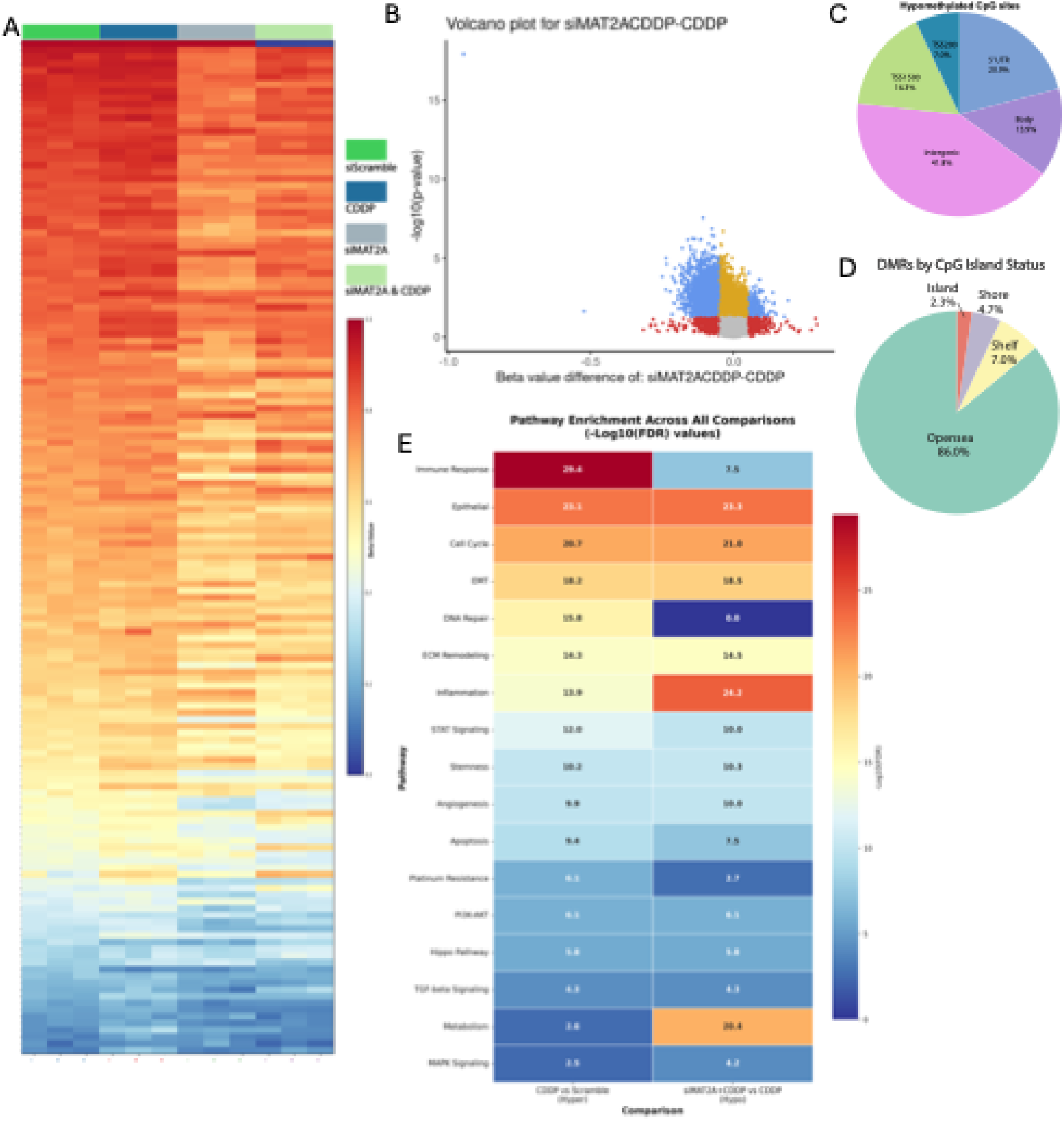
MAT2A inhibition abrogates platinum-induced hypermethylation and reveals extensive TF-driven pathway enrichment. **(A)** Heatmap of beta values for statistically significant DMPs identified across platinum (cisplatin, CDDP; 15μM,16 hours), siMAT2A (48 hours), and siMAT2A + CDDP conditions. **(B)** Volcano plot of CpG-level methylation changes between siMAT2A + CDDP and CDDP-only treated cells. **(C)** Pie chart of the distribution of hypomethylated DMPs across gene features (promoter, exon, intron, and intergenic regions) in siMAT2A + CDDP versus platinum-only treatment. **(D)** Pie chart of genomic annotation of DMRs across CpG island contextual categories (island, shore, shelf, and open sea) in siMAT2A + CDDP versus platinum-only treatment. **(E)** Predicted transcription factor binding site (TFBS)-enriched pathways from two comparisons: hypermethylated DMRs in CDDP versus siScramble (left) and hypomethylated DMRs in siMAT2A + CDDP versus CDDP (right). (* p<0.05, ** p<0.01, *** p<0.001, **** p<0.0001; ns, not significant)

As it is well established that DNA methylation at regulatory regions influences accessibility and binding of transcription factors^59^, probabilistic genomic concordance analysis was performed to predict transcription factor binding sites (TFBS)^41,60^ on hypermethylated DMRs (platinum vs siScramble) and hypomethylated DMRs (platinum+siMAT2A vs platinum). We mapped the corresponding TFs to validated target genes for pathway enrichment, identifying processes potentially affected by methylation changes at TF binding sites. Surprisingly, TFBS analysis revealed several overlapping pathways between hyper- and hypo-methylation changes [Fig.3E], indicating that siMAT2A modulates the TFBS and pathways also regulated by platinum-induced hypermethylation. The predicted-TFBS enriched pathways also revealed that siMAT2A+platinum reduced enrichment of platinum resistance, DNA repair, and immune response pathways and increased enrichment of inflammatory and metabolic pathways [Fig.3E].

### MAT2A inhibition increases platinum sensitivity by altering the DNA damage response (DDR)

Given that platinum acts by inducing DNA damage^61^, the reduced enrichment at TFBS related to DNA repair and platinum resistance pathways [Fig.3E] prompted us to hypothesize that MAT2A inhibition enhances platinum efficiency in multiple HGSC cells. To investigate the role of MAT2A in platinum sensitivity, we assessed apoptosis by AnnexinV-PI staining with or without siMAT2A. Compared to platinum alone, siMAT2A+platinum significantly increased total apoptosis, evidenced by elevated early and late apoptotic fractions [Fig.4A-B; Supplemental Fig.S3A-B]. To test whether the observed increase in sensitivity resulted from enhanced DNA damage, we used the comet to detect DNA breaks and assess migration of fragmented DNA or “tail moment” at the single-cell level. Compared to either platinum or siMAT2A alone, cells treated with siMAT2A+platinum significantly increased tail moments [Fig.4C-D; Supplemental Fig.S3C-D]. MAT2A inhibition can promote R-loop (RNA:DNA hybrids) accumulation, leading to replication stress^18^, we examined whether concurrent R-loop accumulation underlies the siMAT2A-induced increase in DNA damage. Given the potential off-target recognition of cellular RNA by the S9.6 antibody, we focused our analysis on nuclear R-loop signals^62^, as these are directly relevant to R-loop-associated genomic DNA damage. Consistent with enhanced DNA damage, the number of R-loops detected in the nucleus after shMAT2A+platinum treatment was greater compared to platinum or shMAT2A alone in Kuramochi [Fig.4E-F], as well as in OVCAR5 [Supplemental Fig.S3E-F]. However, despite inducing more DNA damage, siMAT2A+platinum unexpectedly decreased γH2AX, compared to platinum alone [Fig.4E,G; Supplemental Fig.S3E,G; Fig.S3I]. In addition to flagging DNA lesions, γH2AX recruits repair proteins^63^ and the reduced γH2AX signal in combination treatment may reflect impaired DDR activation, which would be consistent with the reduced enrichment of the DNA repair pathway identified by the predicted-TFBS pathway analysis of siMAT2A+platinum group [Fig.3E].

**Figure 4.**
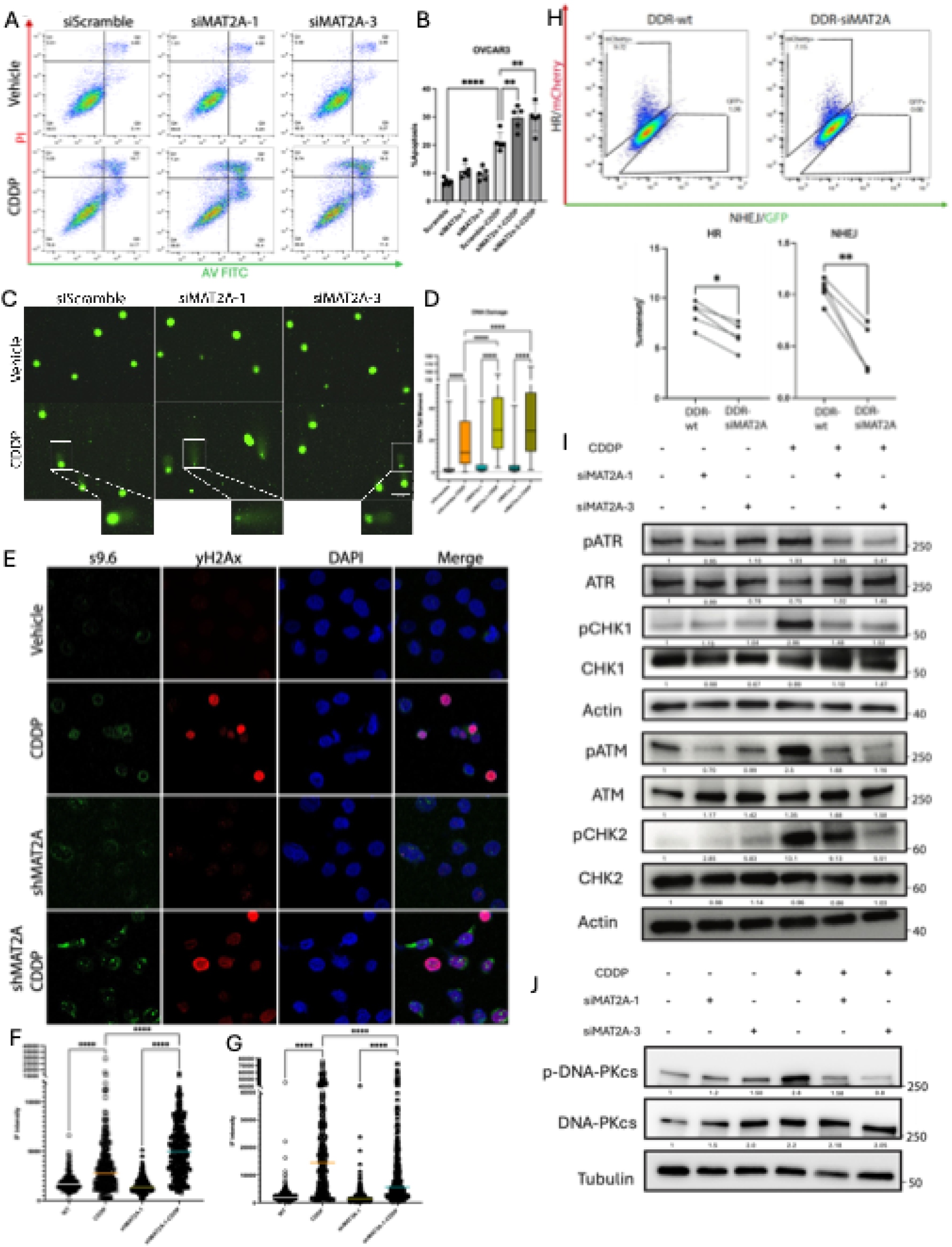
Inhibiting MAT2A increases chemotherapy sensitivity by enhancing DNA damage and reducing DNA repair. **(A-B)** Representative apoptosis assay plots (A) and quantification (B) of %apoptotic cells in OVCAR3 cells treated with siMAT2A-1 (48 hours), siMAT2A-3 (48 hours), platinum (cisplatin, CDDP; 15μM,16 hours), or combination. **(C-D)** Representative comet assay images (C) and quantification (D) of tail moment as a measure of DNA damage in OVCAR3 cells treated with siMAT2A-1, siMAT2A-3, CDDP, or combination. Box plots summarize the distribution of tail moment values for each treatment group; the center line indicates the median, the box represents the interquartile range, and the whiskers indicate the minimum and maximum values. **(E-G)** Representative immunofluorescence images (E) and quantification of S9.6 (F) and γH2AX (G) in Kuramochi wild-type and shMAT2A cells with or without CDDP (12μM,16 hours) treatment. Each dot represents the fluorescence intensity of a single cell, and the horizontal colored line indicates the mean. **(H)** Representative scatter plots of HR-mCherry and NHEJ-GFP reporter activity in OVCAR3-DDR cells with or without siMAT2A treatment, and quantification of HR and NHEJ activity. Each dot represents an independent biological replicate, and lines denote paired measurements obtained from the same biological replicate. **(I-J)** Western blot of key HR (I) and NHEJ (J) pathway proteins (phosphorylated and total) in OVCAR3 under indicated treatment conditions, with densitometric quantification relative to actin or tubulin from the corresponding sample. (* p<0.05, ** p<0.01, *** p<0.001, **** p<0.0001; ns, not significant)

Based on the DNA damage results above, we used a DNA damage reporter system^44^ (DDRS), which simultaneously measures homologous recombination (HR) and non-homologous end joining (NHEJ) repair marked by mCherry or GFP, respectively^44^, at single cell level, to directly tested the effect of MAT2A knockdown on DNA repair. Compared to vector control, MAT2A knockdown significantly decreased both %GFP and %mCherry in HGSC cell lines [Fig.4H; Supplemental Fig.S3J,K], indicating inhibition of both NHEJ and HR. We next examined key HR (ATR, ATM, CHK1, CHK2) and NHEJ (DNA-PKcs, Ku70, Ku80) sensors and transducers using western blot analysis. Similar alterations in both pathways were observed across all treatments [Fig.4I-J; Supplemental Fig.S3L-N]. Unexpectedly, neither platinum nor siMAT2A affected the total protein expression of these sensors and transducers. However, platinum-induced phosphorylation of ATR, CHK1, ATM, CHK2, and DNA-PKcs was abrogated by combining siMAT2A [Fig.4I-J; Supplemental Fig.S3L-N]. Collectively, these data established a novel role of MAT2A in mediating platinum-induced DDR and MAT2A as a target to sensitize HGSC cells to platinum.

### SAM synthesis mediates DDR through a SAMTOR-mTOR-S6K-FANCD2 axis

Having shown that inhibiting MAT2A augmented platinum-induced DDR, we mechanistically examined how HGSC cells abrogate DNA repair activation in response to decreased MAT2A by focusing on signaling pathways downstream of SAM: mTOR signaling, the primary nutrient-sensing pathway for amino acids and p-S6K, a key indicator of mTOR activity^64^. We conducted western blot analysis after MAT2A inhibition and/or platinum treatment. While total mTOR level was not altered by platinum, both siMAT2A and FIDAS-5 inhibited platinum-induced phosphorylation of S6K [Supplemental Fig.S4A,B]. Regulation of mTOR signaling through sensing the level of SAM by S-adenosylmethionine sensor upstream of mTORC1 (SAMTOR, BMT2) has been described^65,66^. When SAM level drops, BMT2 directly interacts with the GATOR1 component Depdc5, inhibiting mTOR activities^65,66^. As indicated by co-IP, the interaction between Flag-Depdc5 and BMT2 was significantly increased by siMAT2A, and SAM supplementation reduced this interaction in a dose-dependent manner [Fig.5A; Supplemental Fig.S4C], demonstrating that knockdown of a metabolic enzyme (MAT2A) can be translated into a cellular response (mTOR signaling) in HGSC.

**Figure 5.**
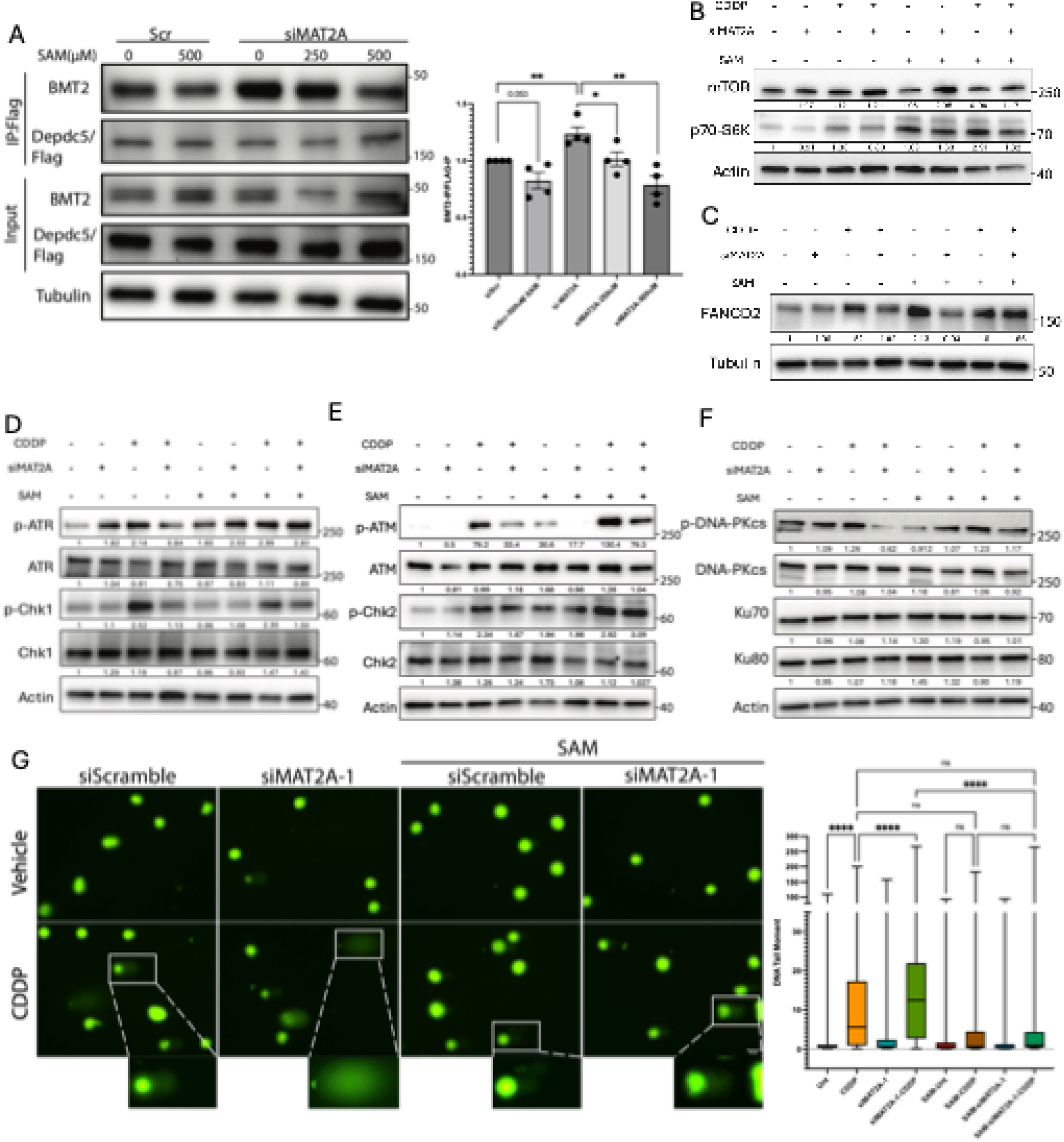
SAM synthesis mediates DNA repair activation in response to platinum treatment through mTOR-S6K-FANCD axis. **(A)** Co-immunoprecipitation (Co-IP) of FLAG-DEPDC5-transfected OVCAR3 cells treated with siMAT2A with SAM concentrations (250 or 500 μM) added at lysate for overnight incubation, and densitometric quantification of BMT2 normalized to Flag-Depdc5. **(B-F)** Western blot of mTOR and p70-S6K (B), FANCD2 (C), HR DNA repair proteins (D-E) and NHEJ DNA repair proteins (F) in OVCAR3 cells treated with platinum (cisplatin, CDDP; 15μM,16 hours), siMAT2A (48 hours), or combination with or without SAM supplementation (50μM, 54 hours), with densitometric quantification normalized to actin. **(G)** Representative comet assay images and quantification of tail moment in OVCAR3 cells treated with siMAT2A (48 hours), CDDP (15μM,16 hours), or combination with or without SAM rescue (50μM, 54 hours). (* p<0.05, ** p<0.01, *** p<0.001, **** p<0.0001; ns, not significant)

Notably, similar responses to MAT2A inhibition were observed among S6K [Supplemental Fig.S4A-B] and multiple DNA damage response proteins [Fig.4I-J; Supplemental Fig.S3L-N], i.e., total protein levels remained largely unchanged, but phosphorylation was markedly reduced, suggesting the possibility that mTOR and DNA damage response proteins may be involved in the same regulatory axis and thus functionally linked. In support of this possibility, Shen et al. reported that mTOR-dependent downregulation of FANC protein attenuates ATM/CHK2 activation^67^, and we observed increased R-loop [Fig.4E-F; Supplemental Fig.S3E-F] and decreased of FANCD2 [Fig.S3H] with siMAT2A+platinum treatment. Together, those results led us to hypothesize that mTOR-S6K-FANCD2 axis mechanistically links SAM synthesis to DNA repair activation. To test this, we examined whether SAM supplementation could rescue the reduced signaling observed after siMAT2A treatment. We first confirmed that the effect of MAT2A knockdown was unaffected by SAM supplementation [Supplementary Fig. S4D-E], indicating that any subsequent rescue effects would be due to SAM rather than changes in MAT2A protein levels. While SAM supplementation largely had no effect on total protein levels across all treatments, increased phosphorylation of p70-S6K, FANCD2, pATR, p-ATM and pCHK2 was observed with SAM supplementation alone [Fig.5B-F; Supplemental Fig.S4F-J]. Adding SAM back to the media reversed the effects of platinum+siMAT2A, restoring FANCD2 expression and partially restoring phosphorylation of S6K, ATM, ATR, CHK1, and CHK2 [Fig.5B-F; Supplemental Fig.S4F-J]. We then used the comet assay to examine whether these altered phosphoprotein levels have a functional impact on DDR. Compared to platinum alone, SAM supplementation together with platinum treatment decreased the comet-like tail [Fig.5G; Supplemental Fig.S4K]. Moreover, when compared to siMAT2A+platinum combination, SAM supplementation reduced tail moments to a level comparable to or lower than platinum alone [Fig.5G; Supplemental Fig.S4K]. Overall, these results indicate that MAT2A-driven SAM synthesis, acting through the SAMTOR-mTOR-S6K-FANCD2 signaling axis, plays a key role in activating DDR to protect HGSC cells from platinum-induced damage.

### SAM synthesis is essential for platinum-enriched OCSCs in both MTAP-proficient and -deficient HGSC

Given that enhanced DDR facilitates OCSC survival after platinum therapy^68^, MAT2A may contribute to OCSC enrichment through DNA repair activation described above. In support of this possibility, IPA analysis of DMRs from the platinum versus siScramble revealed multiple pathways related to stemness characteristics [Fig.2H], a phenotype that requires methyl groups provided by SAM^11^. Consistent with the IPA results, IHC of xenograft tumors demonstrated alterations ALDH1A1 and SIRT1 expression following platinum treatment compared to vehicle [Supplementary Fig.S5A]. Dual nuclear and cytoplasmic staining of SIRT1 was observed following platinum treatment [Supplementary Fig.S5A, right], and this dual localization has been associated with acquired chemoresistance and stemness in OC^69,70^. Together with the platinum-induced increase in MAT2A *in vivo* [Fig.1E], we hypothesized that by regulating DDR and altering the methylome, MAT2A-driven SAM synthesis is essential for platinum-induced enrichment of OCSC.

To investigate whether the Met cycle is required for platinum-induced stemness, HGSC cells were cultured in Met depletion media [Met(-)] and/or treated with the IC_50_ of platinum. Compared to the untreated group, platinum treatment increased the %ALDH+ cells by 1.5-2 fold above untreated baseline [Supplemental Fig.S5B,D] and increased the number of spheroids [Supplemental Fig.S5C,E]. Strikingly, the absence of Met in the media prevented the platinum-induced increase in %ALDH+ cells and spheroid formation regardless of MTAP status [MTAP-high OVCAR3, Fig.S5B,C; MTAP-null OVCAR5, Supplemental Fig.S5D,E]. We supplemented HGSC cells with exogenous SAM and at all concentrations tested [25-200µM of SAM], in the absence of platinum treatment and regardless of MTAP status, SAM increased the %ALDH+ cells [Fig.6A; Supplemental Fig.S5F]. Taken together, these results support the critical role of Met cycle in OCSC.

**Figure 6.**
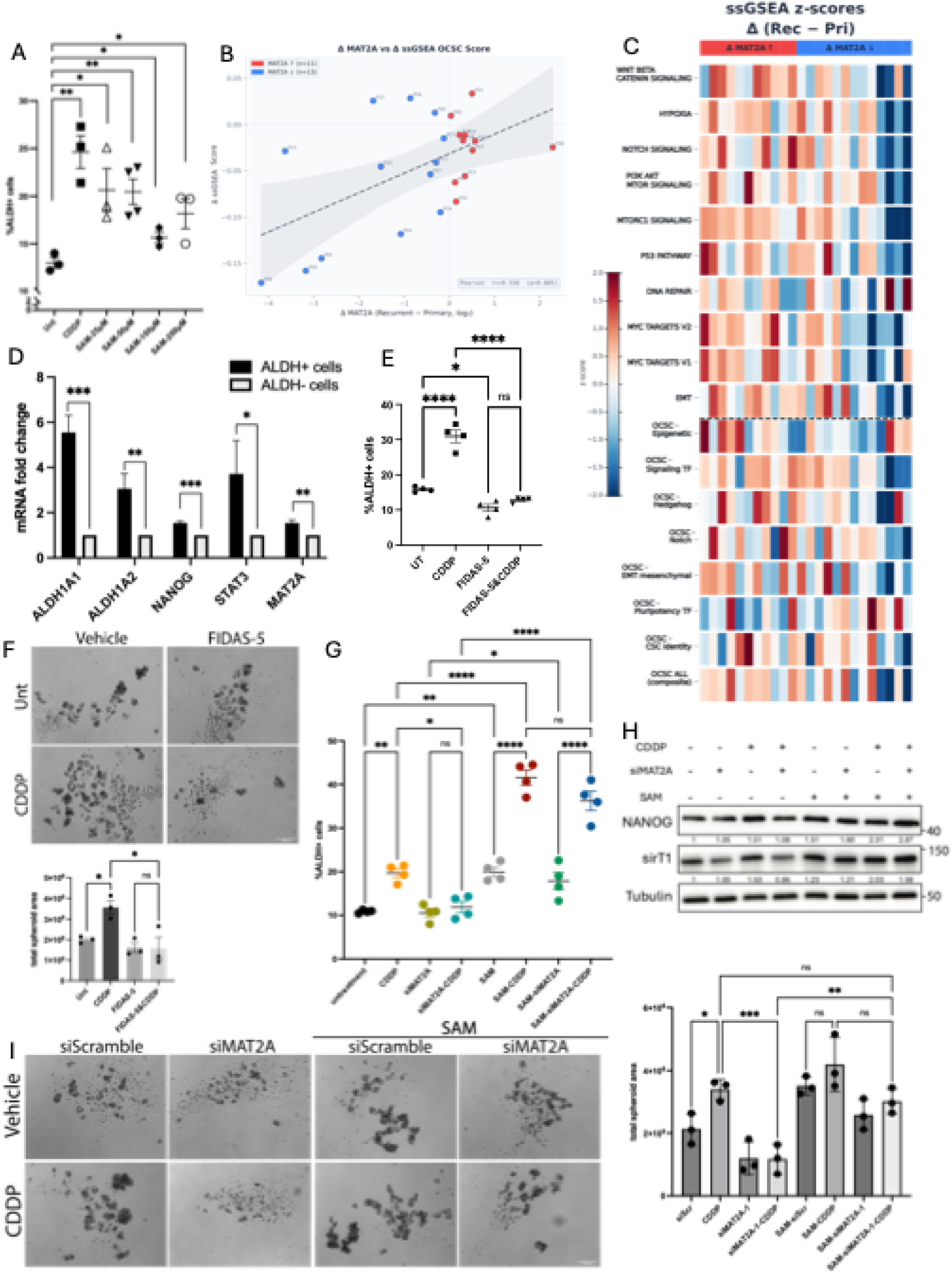
MAT2A and SAM are essential for platinum-induced OCSC phenotype and recurrent OC. **(A)** Quantification of percentage of ALDH+ cells in OVCAR3 cells supplemented with platinum (cisplatin, CDDP; 15μM,16 hours) or increasing concentrations of exogenous SAM (25-200μM, 54 hours). **(B)** ssGSEA-derived OCSC scores plotted against ΔMAT2A expression between matched recurrent and primary tumors. **(C)** Heatmap of ssGSEA z-scores for OCSC markers and validated CSC hallmark gene sets in patients stratified by ΔMAT2A direction (upregulated and downregulated in recurrent relative to primary tumors). **(D)** Relative mRNA expression of MAT2A and stemness markers in ALDH+ and ALDH− cell fractions isolated by FACS. **(E-F)** Percentage of ALDH+ cell (E), and representative images and quantification of spheroid formation (F) in OVCAR3 cells treated with FIDAS-5 (11.8μM, 48 hours), CDDP (15μM, 16 hours) or combination. **(G)** Percentage of ALDH+ cells in OVCAR3 cells treated with CDDP (15μM,16 hours), siMAT2A (48 hours), or combination with or without 50 μM exogenous SAM (54 hours). **(H)** Stemness marker protein expression by western blot with densitometric quantification to actin or tubulin in OVCAR3 cells treated with CDDP (15μM,16 hours), siMAT2A (48 hours), or combination with or without 50 μM exogenous SAM (54 hours). **(I)** Representative images and quantification of spheroid formation in OVCAR3 cells treated with CDDP (15μM,16 hours), siMAT2A (48 hours), or combination with or without 50 μM exogenous SAM (54 hours). (* p<0.05, ** p<0.01, *** p<0.001, **** p<0.0001; ns, not significant)

To assess whether MAT2A upregulation in recurrent HGSC tumors was coupled to transcriptional reprogramming to OCSC, ssGSEA analysis was performed on 24 paired primary and recurrent (post-chemotherapy) tumor samples from patients diagnosed with HGSC^8^. The composite CSC ssGSEA score integrates enrichment across seven functionally defined CSC subcategories curated from the literature [Supplemental Table S2]. Patient HGSC tumors with increased MAT2A expression at recurrence exhibited a corresponding increase in OCSC composite score [Fig.6B; Supplemental Fig. S5G], indicated by a positive association (Pearson r=+0.556, p=0.005). The paired ΔssGSEA heatmap, incorporating ten MSigDB Hallmark sets^35,36^ as external validation, revealed consistent enrichment of stemness-associated hallmarks in tumors with elevated MAT2A expression from patients with recurrent HGSC [Fig.6C]. Furthermore, higher expression of MAT2A mRNA together with selected stemness markers were observed in ALDH+ vs. ALDH-cells [Fig.6D].

Based on these results, we examined whether inhibiting MAT2A pharmacologically with FIDAS-5 would suppresses OCSC functional properties in vitro. Treatment of HGSC cell lines with FIDAS-5 blocked the platinum-induced increase in %ALDH+ cells and decreased spheroid formation ability [Fig.6E,F; Supplemental Fig.S5H,I]. Genetically knocking down MAT2A achieved essentially similar effects in all HGSC cell lines tested [Supplemental Fig.S6A-E]. SAM supplementation enhanced the platinum-induced increase in %ALDH+ cells, beyond platinum or SAM alone [Fig.6G; Supplemental Fig.S6F]. Furthermore, in the presence of siMAT2A, exogenous SAM restored the platinum-induced increase in the ALDH+ population [Fig.6G; Supplemental Fig.S6F] and rescued the reduction in stemness markers and spheroid formation [Fig.6H,I; Supplemental Fig.S6G,H]. Importantly, as essentially similar results were seen in MTAP-proficient (OVCAR3) and -deficient (OVCAR5) cells, the data suggest that stemness regulation by MAT2A is mediated through SAM synthesis and not by synthetic lethality with MTAP depletion.

## Discussion

Platinum resistance and disease recurrence remain formidable challenges in HGSC^6^, and a better understanding of how cancer cells adapt to platinum-induced cytotoxic stress and survive is essential to prevent disease relapse and recurrence. Altered metabolic pathways in platinum-resistant HGSC include the Met cycle, which generates the universal methyl donor SAM, an essential metabolic/epigenetic checkpoint linking nutrient sensing to epigenetic regulation^9,31^. Here, we demonstrate that MAT2A-driven SAM synthesis protects HGSC cells from platinum-induced cytotoxic stress through coordinated epigenetic remodeling, DNA damage response, and OCSC enrichment, which has been shown to play a critical role in disease relapse and platinum resistance in HGSC and other cancers^24,27^.

Our findings have several important implications. First, beyond the association between high MAT2A expression and poor PFS, the negative correlation between post-NACT MAT2A expression and patient outcome suggests that MAT2A could serve as a prognostic biomarker to identify patients at high risk of early recurrence after NACT and those who may benefit from MAT2A-targeted intervention. Unlike the selective synthetic lethality between MAT2A inhibition and MTAP deletion reported in several cancer types^17,18^, the present findings demonstrate that MAT2A targeting is effective in both MTAP-proficient and -deficient HGSC cell lines, supporting a broader therapeutic potential for MAT2A inhibition beyond MTAP-deficient tumors.

Second, by DNA methylation profiling analysis, we show that platinum is an “epigenetic stressor” and reshapes the HGSC cells DNA methylome. Our findings support previous studies demonstrating that promoter hypermethylation is a hallmark of platinum-resistant HGSC^71,72^ and extend those observations by directly demonstrating that platinum treatment itself broadly alters the methylome before the establishment of platinum resistance [Fig.2]. We further show MAT2A-driven SAM synthesis is essential for platinum to induce remodeling of the HGSC methylome, highlighted by the ability of siMAT2A to abrogate platinum-induced alterations and subsequent cellular responses. Interestingly, upregulation of MAT2A in recurrent, platinum resistant HGSC tumors appear to be coordinated with an increase in DNMTs, further supporting co-adaptation of metabolic and epigenetic machinery under the pressure of cytotoxic chemotherapy.

Third, by using DNA methylation profiling to build on epigenetic alterations, this study reveals a previously unknown link between MAT2A-driven SAM synthesis and DNA repair in HGSC. A similar phenomena has been reported in glioblastoma^73^ and acute myeloid leukaemia^74^, yet we demonstrate that the linkage is a comprehensive mechanism in HGSC, showing for the first time that MAT2A-driven SAM synthesis mediates the activation of DNA damage response to platinum. While inhibiting MAT2A alters expression of repair proteins by regulating histone methylation^73,74^, this is the first report to quantitatively show a direct reduction in activities of two primary DNA double-strand break repair pathways in living cells following MAT2A knockdown. Mechanistically, MAT2A influences phosphorylation of DNA repair Rescue of this effect of MAT2A knockdown by using exogenous SAM further supports the role of SAM in maintaining DNA repair under platinum stress and extends the concept of “SAM checkpoint” in pre-replication complex and replication stress^75^. These findings overall provide a better understanding and possible explanation for why interrupting the Met cycle enhances chemotherapy and irradiation sensitivity^76,77^.

Finally, we show that MAT2A-driven SAM synthesis not only mediates DNA damage response but also plays a key role in OCSC reprograming. Our bioinformatic analysis of platinum-induced methylomic changes shows striking enrichment of stemness pathways, advancing the concept of an epigenetic contribution to CSC and advancing the concept put forth by us and others that a critical consequence of platinum-based chemotherapy is the conversion of non-CSCs to CSCs^6,38,45,78^. In addition, coupling of post-chemotherapy MAT2A upregulation with CSC gene signature enrichment in patients with recurrent HGSC provides strong support in an epigenetic-metabolism context: MAT2A-driven SAM synthesis is required for OCSC enrichment following platinum treatment. Based on the findings of this study, we suggest that altered MAT2A-SAM metabolism after chemotherapy enhances DNA repair and expands the CSC population, contributing to platinum resistance and ultimately disease recurrence in HGSC patients.

While our findings are based on the use of appropriate in vitro models and robust bioinformatic analysis of tumors from patients in important databases, we recognize that to fully establish the clinical relevance of our pre-clinical findings will require additional model systems such as xenografts or patient-derived organoids, which we will pursue following optimization of pharmacologic and genetic approaches in both MTAP-proficient and -deficient HGSC. Pathway enrichment based on predicted-TFBS demonstrates the impact of methylome dynamics but lacks the resolution to show direction (activated or suppressed). We acknowledge that the downstream transcriptional consequences of platinum-induced epigenetic reprogramming remain to be fully defined. We also recognize that shifting the production of SAM will have an impact beyond DNA methylation including protein and histone methylation, and reduction in histone methylation and DNA methylation may contribute to the increase in R-Loop formation observed in this study. Furthermore, to define methylome changes causally linked to resistance phenotypes would require additional approaches such as multi-omics. We note that other approaches such as lineage tracing would be necessary to fully understand how MAT2A-driven SAM synthesis reprograms cells that survive platinum treatment toward a stem-like phenotype and/or supports existing OCSCs.

In summary, we show that MAT2A plays key roles both in platinum-induced epigenetic adaptation and DDR in HGSC. Targeting MAT2A could inhibit metabolic-epigenetic dependencies, block post-chemotherapy enrichment of OCSC and therefore represent a therapeutic strategy to resensitize tumors to platinum.

## Acknowledgment

We would like to thank the VAI Bioinformatics and Biostatistics Core for their assistance with profiling of EpicArray samples, and Sheldon R. and Metabolomics & Bioenergetics Core for their assistance of LC-MS analysis. We also thank the Flow Cytometry Core Facility and Light Microscopy Imaging Center for their technical support, and the Histology Lab Service Core and Laboratory Animal Resource Center for assistance with the mouse study.

This research was supported in part by the Ovarian Cancer Alliance of Greater Cincinnati (to K.P. Nephew and H.M. O’Hagan), the IUSCCC P30 Support Grant by Indiana University (NCI Grant Number P30CA082709-25), the Congressionally Directed Medical Research Programs-Ovarian Cancer Research Program (to K.P. Nephew and H.M. O’Hagan), Department of Defense Ovarian Cancer Research Program Grant Number W81XWH-21-1-0284, and Van Andel Institute through the Van Andel Institute - Stand Up To Cancer Epigenetics Dream Team. We also express our gratitude to the Jerry and Peggy Throgmartin Family for their philanthropic investments in our work. Shu Zhang was supported by the Doane and Eunice Dahl Wright Fellowship generously provided by Ms. Imogen Dahl.

## CRediT authorship contribution statement

**Shu Zhang:** Conceptualization, Data curation, Formal analysis, Methodology, Software, Validation, Visualization, Writing - original draft, Writing - review & editing. **Zhen Fu:** Data curation, Methodology, Software, Visualization. **Yanchi Zhou:** Conceptualization, Data curation, Methodology, Writing - review & editing. **Tara X. Metcalfe:** Conceptualization, Data curation, Methodology, Writing - review & editing. **Truc T. Vuong:** Data curation, Methodology. **Lindsay W. Brubaker:** Methodology, Writing - review & editing. **Benjamin G. Bitler:** Methodology, Visualization, Writing - review & editing. **Heather M. O’Hagan:** Conceptualization, Data curation, Methodology, Visualization, Writing - original draft, Writing - review & editing. **Kenneth P. Nephew:** Conceptualization, Formal analysis, Funding acquisition, Investigation, Methodology, Project administration, Resources, Visualization, Writing - original draft, Writing - review & editing.

## References

1 Kurnit, K. C., Fleming, G. F. & Lengyel, E. Updates and New Options in Advanced Epithelial Ovarian Cancer Treatment. Obstet Gynecol 137, 108–121 (2021). 10.1097/AOG.0000000000004173

2 Mahmood, R. D., Morgan, R. D., Edmondson, R. J., Clamp, A. R. & Jayson, G. C. First-Line Management of Advanced High-Grade Serous Ovarian Cancer. Curr Oncol Rep 22, 64 (2020). 10.1007/s11912-020-00933-8

3 Berek, J. S. et al. Advanced epithelial ovarian cancer: 1998 consensus statements. Ann Oncol 10 **Suppl 1**, 87–92 (1999). 10.1023/a:1008323922057

4 Vaughan, S. et al. Rethinking ovarian cancer: recommendations for improving outcomes. Nat Rev Cancer 11, 719–725 (2011). 10.1038/nrc3144

5 Javellana, M. et al. Neoadjuvant Chemotherapy Induces Genomic and Transcriptomic Changes in Ovarian Cancer. Cancer Res 82, 169–176 (2022). 10.1158/0008-5472.CAN-21-1467

6 Balkwill, F. R. et al. Rethinking ovarian cancer III: the past decade and future directions. Nat Rev Cancer 26, 452–471 (2026). 10.1038/s41568-026-00916-0

7 Rizzo, A. et al. One-Carbon Metabolism: Biological Players in Epithelial Ovarian Cancer. Int J Mol Sci 19 (2018). 10.3390/ijms19072092

8 Kim, M. A. et al. Integrative Multi-Omics Analysis Reveals Molecular Signatures of Recurrence in Paired Primary and Recurrent High-Grade Serous Ovarian Cancer. Int J Mol Sci 27 (2026). 10.3390/ijms27020948

9 Poisson, L. M. et al. A metabolomic approach to identifying platinum resistance in ovarian cancer. J Ovarian Res 8, 13 (2015). 10.1186/s13048-015-0140-8

10 Hoffman, R. M. & Erbe, R. W. High in vivo rates of methionine biosynthesis in transformed human and malignant rat cells auxotrophic for methionine. Proc Natl Acad Sci U S A 73, 1523–1527 (1976). 10.1073/pnas.73.5.1523

11 Wang, B. Q. et al. Unraveling the potential of targeting methionine metabolism in cancer. Cancer Lett 644, 218338 (2026). 10.1016/j.canlet.2026.218338

12 Han, Q., Tan, Y. & Hoffman, R. M. Oral dosing of Recombinant Methioninase Is Associated With a 70% Drop in PSA in a Patient With Bone-metastatic Prostate Cancer and 50% Reduction in Circulating Methionine in a High-stage Ovarian Cancer Patient. Anticancer Res 40, 2813–2819 (2020). 10.21873/anticanres.14254

13 Capellini, E. et al. Intermittent fasting enhances cisplatin-metformin efficacy in therapy-resistant ovarian cancer PDXs. iScience 29, 114300 (2026). 10.1016/j.isci.2025.114300

14 Tian, X., Zhu, G., Zhang, Y. & Liu, N. Methionine restriction in cancer: a dietary insight for therapy. Front Nutr 13, 1730639 (2026). 10.3389/fnut.2026.1730639

15 Kawaguchi, K. et al. Efficacy of Recombinant Methioninase (rMETase) on Recalcitrant Cancer Patient-Derived Orthotopic Xenograft (PDOX) Mouse Models: A Review. Cells 8 (2019). 10.3390/cells8050410

16 Bandaru, N. et al. Methionine restriction for cancer therapy: From preclinical studies to clinical trials. Cancer Pathog Ther 4, 124–135 (2026). 10.1016/j.cpt.2025.01.002

17 Bedard, G. T. et al. Combined inhibition of MTAP and MAT2a mimics synthetic lethality in tumor models via PRMT5 inhibition. J Biol Chem 300, 105492 (2024). 10.1016/j.jbc.2023.105492

18 Kalev, P. et al. MAT2A Inhibition Blocks the Growth of MTAP-Deleted Cancer Cells by Reducing PRMT5-Dependent mRNA Splicing and Inducing DNA Damage. Cancer Cell 39, 209–224 e211 (2021). 10.1016/j.ccell.2020.12.010

19 Rodon, J., Johnson, M. L., George, B., Shah, P. A. & Arbour, K. C. MTAP Deletion in Oncogenesis: A Synthetic Lethality Scenario. Cancer Res 86, 1558–1569 (2026). 10.1158/0008-5472.CAN-25-2126

20 Zhu, J. et al. Transsulfuration Activity Can Support Cell Growth upon Extracellular Cysteine Limitation. Cell Metab 30, 865–876 e865 (2019). 10.1016/j.cmet.2019.09.009

21 Traverso, N. et al. Role of glutathione in cancer progression and chemoresistance. Oxid Med Cell Longev 2013, 972913 (2013). 10.1155/2013/972913

22 Zhang, S. et al. Identification and characterization of ovarian cancer-initiating cells from primary human tumors. Cancer Res 68, 4311–4320 (2008). 10.1158/0008-5472.CAN-08-0364

23 Bapat, S. A., Mali, A. M., Koppikar, C. B. & Kurrey, N. K. Stem and progenitor-like cells contribute to the aggressive behavior of human epithelial ovarian cancer. Cancer Res 65, 3025–3029 (2005). 10.1158/0008-5472.CAN-04-3931

24 Zong, X. & Nephew, K. P. Ovarian Cancer Stem Cells: Role in Metastasis and Opportunity for Therapeutic Targeting. Cancers (Basel) 11 (2019). 10.3390/cancers11070934

25 Manzoor, H. B. et al. Global DNA methylation signatures associated with chemoresistance and poor prognosis of high grade serous ovarian cancer. Sci Rep 15, 36869 (2025). 10.1038/s41598-025-20827-8

26 Sriramkumar, S. et al. Platinum-induced mitochondrial OXPHOS contributes to cancer stem cell enrichment in ovarian cancer. J Transl Med 20, 246 (2022). 10.1186/s12967-022-03447-y

27 Balch, C., Fang, F., Matei, D. E., Huang, T. H. & Nephew, K. P. Minireview: epigenetic changes in ovarian cancer. Endocrinology 150, 4003–4011 (2009). 10.1210/en.2009-0404

28 Zeller, C. et al. Candidate DNA methylation drivers of acquired cisplatin resistance in ovarian cancer identified by methylome and expression profiling. Oncogene 31, 4567–4576 (2012). 10.1038/onc.2011.611

29 Fang, F. et al. A phase 1 and pharmacodynamic study of decitabine in combination with carboplatin in patients with recurrent, platinum-resistant, epithelial ovarian cancer. Cancer 116, 4043–4053 (2010). 10.1002/cncr.25204

30 Vuong, T. T. et al. DNA Methyltransferase Inhibition Prevents Platinum-Induced Ovarian Cancer Stem Cell Enrichment. Cancer Research Communications (2026). 10.1158/2767-9764.Crc-26-0149

31 Wang, Z. et al. Methionine is a metabolic dependency of tumor-initiating cells. Nat Med 25, 825–837 (2019). 10.1038/s41591-019-0423-5

32 Strekalova, E. et al. S-adenosylmethionine biosynthesis is a targetable metabolic vulnerability of cancer stem cells. Breast Cancer Res Treat 175, 39–50 (2019). 10.1007/s10549-019-05146-7

33 Haley, J. et al. Functional characterization of a panel of high-grade serous ovarian cancer cell lines as representative experimental models of the disease. Oncotarget 7, 32810–32820 (2016). 10.18632/oncotarget.9053

34 Zhang, H. et al. Integrated Proteogenomic Characterization of Human High-Grade Serous Ovarian Cancer. Cell 166, 755–765 (2016). 10.1016/j.cell.2016.05.069

35 Fang, Z., Liu, X. & Peltz, G. GSEApy: a comprehensive package for performing gene set enrichment analysis in Python. Bioinformatics 39 (2023). 10.1093/bioinformatics/btac757

36 Liberzon, A. et al. The Molecular Signatures Database (MSigDB) hallmark gene set collection. Cell Syst 1, 417–425 (2015). 10.1016/j.cels.2015.12.004

37 Virtanen, P. et al. SciPy 1.0: fundamental algorithms for scientific computing in Python. Nat Methods 17, 261–272 (2020). 10.1038/s41592-019-0686-2

38 Wang, W. et al. Targeting Ovarian Cancer Stem Cells by Dual Inhibition of the Long Noncoding RNA HOTAIR and Lysine Methyltransferase EZH2. Mol Cancer Ther 23, 1666–1679 (2024). 10.1158/1535-7163.MCT-23-0314

39 Noguera-Castells, A., Garcia-Prieto, C. A., Alvarez-Errico, D. & Esteller, M. Validation of the new EPIC DNA methylation microarray (900K EPIC v2) for high-throughput profiling of the human DNA methylome. Epigenetics 18, 2185742 (2023). 10.1080/15592294.2023.2185742

40 Zhou, W., Triche, T. J., Jr., Laird, P. W. & Shen, H. SeSAMe: reducing artifactual detection of DNA methylation by Infinium BeadChips in genomic deletions. Nucleic Acids Res 46, e123 (2018). 10.1093/nar/gky691

41 Han, H. et al. TRRUST v2: an expanded reference database of human and mouse transcriptional regulatory interactions. Nucleic Acids Res 46, D380–D386 (2018). 10.1093/nar/gkx1013

42 Maurano T. M., W. H., John S., Shafer A., Canfield T., Lee K., Stamatoyannopoulos A. J. Role of DNA Methylation in Modulating Transcription Factor Occupancy,. Cell Reports 12, 1184–1195 (2015). 10.1016/j.celrep.2015.07.024.

43 Jones, P. A. Functions of DNA methylation: islands, start sites, gene bodies and beyond. Nat Rev Genet 13, 484–492 (2012). 10.1038/nrg3230

44 Arnoult, N. et al. Regulation of DNA repair pathway choice in S and G2 phases by the NHEJ inhibitor CYREN. Nature 549, 548–552 (2017). 10.1038/nature24023

45 Zong, X. et al. EZH2-Mediated Downregulation of the Tumor Suppressor DAB2IP Maintains Ovarian Cancer Stem Cells. Cancer Res 80, 4371–4385 (2020). 10.1158/0008-5472.CAN-20-0458

46 Acland, M. et al. Chemoresistant Cancer Cell Lines Are Characterized by Migratory, Amino Acid Metabolism, Protein Catabolism and IFN1 Signalling Perturbations. Cancers (Basel) 14 (2022). 10.3390/cancers14112763

47 Casero, R. A., Jr., Murray Stewart, T. & Pegg, A. E. Polyamine metabolism and cancer: treatments, challenges and opportunities. Nat Rev Cancer 18, 681–695 (2018). 10.1038/s41568-018-0050-3

48 Gyorffy, B. Discovery and ranking of the most robust prognostic biomarkers in serous ovarian cancer. Geroscience 45, 1889–1898 (2023). 10.1007/s11357-023-00742-4

49 Chang, T. Y. et al. Integrated Spatial Analysis Reveals the Molecular Landscape of Ovarian Precancerous Lesions. Cancer Res 86, 1739–1752 (2026). 10.1158/0008-5472.CAN-25-3189

50 Hope A. Townsend, K. R. J., Rebecca J. Wolsky, Lucy B. Van Kleunen, Natalie R. Davidson, Kian Behbakht, Matthew J. Sikora, Robin D. Dowell, Aaron Clauset, Benjamin G. Bitler. Cross-assay RNA modeling reveals cancer biomarkers. (2026). 10.64898/2026.04.30.722009

51 Adzibolosu, N. et al. Immunological modifications following chemotherapy are associated with delayed recurrence of ovarian cancer. Front Immunol 14, 1204148 (2023). 10.3389/fimmu.2023.1204148

52 Ikushima, H., Watanabe, K., Shinozaki-Ushiku, A., Oda, K. & Kage, H. Pan-cancer clinical and molecular landscape of MTAP deletion in nationwide and international comprehensive genomic data. ESMO Open 10, 104535 (2025). 10.1016/j.esmoop.2025.104535

53 Nilforoushan, N. & Moatamed, N. A. Evaluation of MTAP immunohistochemistry loss of expression in ovarian serous borderline tumors as a potential marker for prognosis and progression. Ann Diagn Pathol 48, 151582 (2020). 10.1016/j.anndiagpath.2020.151582

54 Jin, H. et al. Systematic transcriptional analysis of human cell lines for gene expression landscape and tumor representation. Nat Commun 14, 5417 (2023). 10.1038/s41467-023-41132-w

55. Atlas, T. H. P. The Human Protein Atlas v24.0, <https://www.proteinatlas.org> (2026).

56 Chan, D. W. et al. Genome-wide DNA methylome analysis identifies methylation signatures associated with survival and drug resistance of ovarian cancers. Clin Epigenetics 13, 142 (2021). 10.1186/s13148-021-01130-5

57 Zivic, K. et al. The Effects of BRCA1 and BRCA2 Promoter Methylation on Clinicopathological Characteristics and Clinical Outcomes in HGSOC. Cells 15 (2026). 10.3390/cells15030277

58 Krupenko, S. A. & Krupenko, N. I. Loss of ALDH1L1 folate enzyme confers a selective metabolic advantage for tumor progression. Chem Biol Interact 302, 149–155 (2019). 10.1016/j.cbi.2019.02.013

59 Guertin, M. J. & Lis, J. T. Mechanisms by which transcription factors gain access to target sequence elements in chromatin. Curr Opin Genet Dev 23, 116–123 (2013). 10.1016/j.gde.2012.11.008

60 Castro-Mondragon, J. A. et al. JASPAR 2022: the 9th release of the open-access database of transcription factor binding profiles. Nucleic Acids Res 50, D165–D173 (2022). 10.1093/nar/gkab1113

61 Basu, A. & Krishnamurthy, S. Cellular responses to Cisplatin-induced DNA damage. J Nucleic Acids 2010 (2010). 10.4061/2010/201367

62 Smolka, J. A., Sanz, L. A., Hartono, S. R. & Chedin, F. Recognition of RNA by the S9.6 antibody creates pervasive artifacts when imaging RNA:DNA hybrids. J Cell Biol 220 (2021). 10.1083/jcb.202004079

63 Podhorecka, M., Skladanowski, A. & Bozko, P. H2AX Phosphorylation: Its Role in DNA Damage Response and Cancer Therapy. J Nucleic Acids 2010 (2010). 10.4061/2010/920161

64 Yue, S., Li, G., He, S. & Li, T. The Central Role of mTORC1 in Amino Acid Sensing. Cancer Res 82, 2964–2974 (2022). 10.1158/0008-5472.CAN-21-4403

65 Tang, X. et al. Molecular mechanism of S-adenosylmethionine sensing by SAMTOR in mTORC1 signaling. Sci Adv 8, eabn3868 (2022). 10.1126/sciadv.abn3868

66 Gu, X. et al. SAMTOR is an S-adenosylmethionine sensor for the mTORC1 pathway. Science 358, 813–818 (2017). 10.1126/science.aao3265

67 Shen, C. et al. Regulation of FANCD2 by the mTOR pathway contributes to the resistance of cancer cells to DNA double-strand breaks. Cancer Res 73, 3393–3401 (2013). 10.1158/0008-5472.CAN-12-4282

68 Alizadeh, H., Akbarabadi, P., Dadfar, A., Tareh, M. R. & Soltani, B. A comprehensive overview of ovarian cancer stem cells: correlation with high recurrence rate, underlying mechanisms, and therapeutic opportunities. Mol Cancer 24, 135 (2025). 10.1186/s12943-025-02345-3

69 Mvunta, D. H. et al. SIRT1 Regulates the Chemoresistance and Invasiveness of Ovarian Carcinoma Cells. Transl Oncol 10, 621–631 (2017). 10.1016/j.tranon.2017.05.005

70 Xu, H. et al. Cytoplasmic SIRT1 enhances the stemness of polyploid giant cancer cells by promoting beta-catenin protein stability and nuclear accumulation in ovarian carcinoma upon neoadjuvant chemotherapy. Cancer Lett 639, 218193 (2026). 10.1016/j.canlet.2025.218193

71 Ho, C. M., Yen, T. L., Chien, T. Y. & Huang, S. H. Distinct promotor methylation at tumor suppressive genes in ovarian cancer stromal progenitor cells and ovarian cancer and its clinical implication. Am J Cancer Res 12, 5325–5341 (2022).

72 Li, M. et al. Integrated analysis of DNA methylation and gene expression reveals specific signaling pathways associated with platinum resistance in ovarian cancer. BMC Med Genomics 2, 34 (2009). 10.1186/1755-8794-2-34

73 Korimerla, N. et al. Reciprocal links between methionine metabolism, DNA repair and therapy resistance in glioblastoma. bioRxiv (2024). 10.1101/2024.11.20.624542

74 Zeka, K. et al. MAT2A inhibition in AML unveils therapeutic potential of combining DNA demethylating agents with UPR targeting. bioRxiv, 2023.2006.2005.543499 (2023). 10.1101/2023.06.05.543499

75 Lauinger, L. & Kaiser, P. Sensing and Signaling of Methionine Metabolism. Metabolites 11 (2021). 10.3390/metabo11020083

76 Zhao, X. et al. Inhibition of MAT2A-Related Methionine Metabolism Enhances The Efficacy of Cisplatin on Cisplatin-Resistant Cells in Lung Cancer. Cell J 24, 204–211 (2022). 10.22074/cellj.2022.7907

77 Koturbash, I. 2017 Michael Fry Award Lecture When DNA is Actually Not a Target: Radiation Epigenetics as a Tool to Understand and Control Cellular Response to Ionizing Radiation. Radiat Res 190, 5–11 (2018). 10.1667/RR15027.1

78 Muralikrishnan, V., Hurley, T. D. & Nephew, K. P. Targeting Aldehyde Dehydrogenases to Eliminate Cancer Stem Cells in Gynecologic Malignancies. Cancers (Basel) 12 (2020). 10.3390/cancers12040961

