## Supplementary Figures S1-S6 for "Targeting the MAT2A-SAM axis increases platinum sensitivity in ovarian cancer by impairing DNA repair and stemness"

**Supplementary Figure S1. OC Methionine dependency and MAT2A inhibition suppress OC tumor growth**

**(A-B)** Proliferation of OVCAR5 (A) and Kuramochi (B) in complete or methionine-depleted [Met(–)] medium over five days. Data are presented as the mean ± SD of three biological replicates.

**(C)** Met cycle flux map in platinum-resistant versus sensitive CaOV3 cells. Bar graphs represent log2 fold-change (resistant/sensitive) for each metabolite, TIC-normalized and clamped at ±2. Statistical comparisons by Welch's t-test.

**(D)** MAT2A protein abundance in normal fallopian tube versus HGSC tumor tissue of paired samples from the CPTAC dataset.

**(E)** MAT2A mRNA expression across stromal tissue, normal fallopian tube epithelium (NFT), p53 signature (p53 sig), serous tubal intraepithelial lesion (STIL), serous tubal intraepithelial carcinoma (STIC), and high-grade serous carcinoma (HGSC), using data from the Ovarian Pre-cancer Atlas. Boxplots show normalized MAT2A gene counts with individual samples overlaid. Statistical comparisons were performed relative to NFT.

**(F)** Experimental design of the OVCAR3 xenograft model. Female NSG mice were subcutaneously injected with 5 × 10^6 OVCAR3 cells on Day 0 and treated with vehicle or cisplatin (2 mg/kg, intraperitoneally, once weekly) from Day 52 to Day 67.

**(G)**Progression-free survival (PFS) stratified by post-NACT MAT2A expression direction across pooled HGSOC cohorts. Mann-Whitney U test, p = 0.077. Dashed line indicates overall median PFS across all patients.

**(H-I)** Dose-response curves for FIDAS-5 in OVCAR3 (F) and OVCAR5 (G) at 24, 48, 72, and 96 hours. Cell viability (%) was assessed by MTT assay and plotted against $\log_{10}$-transformed FIDAS-5 concentration (μM). IC50 values were derived from four-parameter logistic curve fitting; data represent mean ± SD of three independent experiments.

**(J)** Proliferation assay of OVCAR5 with platinum (cisplatin, CDDP; 3 hours 7.5μM) and/or FIDAS-5 (48hr, 10.9μM) treatment over five days. Data are presented as the mean ± SD of three biological replicates. Symbols (*, #, †) indicate statistical significance ($p$<0.05) for the following comparisons: *, Unt vs. other treatments; #, CDDP vs. FIDAS-5 or combination; †, FIDAS-5 vs. combination. Higher symbol densities (e.g., **, ***) represent $p$<0.01 and $p$ <0.001, respectively.

**Supplementary Figure S2. MAT2A siRNA validation and MAT2A inhibition abrogate platinum-induced hypermethylation**

**(A)** MAT2A mRNA expression in cells in OVCAR5 (left) and OVCAR3 (right) treated with scrambled RNA (siScr) control and MAT2A siRNAs.

**(B)** MAT2A protein level in OVCAR3 treated with scrambled RNA (siScr) control and MAT2A siRNAs.

**(C)** Methylation-specific PCR (MSP) of BRCA1 promoter methylation level. (left) Validation of primers for the BRCA1 (left) promoters using DNA standards with methylation levels ranging from 0% (unmethylated) to 100% (fully methylated), demonstrating clear discrimination between unmethylated- and methylated-specific primer amplification. (middle&right) BRCA1 promoter methylation levels in OVCAR3 cells treated with platinum (cisplatin, CDDP; 15μM, 16 hours), FIDAS-5 (11.8μM, 48 hours)/siMAT2A(48 hours), or FIDAS-5/siMAT2A + CDDP.

**(D)** Methylation-specific PCR (MSP) of ALDH1L1 promoter methylation level. (left) Validation of primers for the ALDH1L1 (left) promoters. (middle & right) ALDH1L1 promoter methylation levels in OVCAR3 cells with indicated treatment.

**(G)** PCA of genome-wide DNA methylation profiles across four treatment conditions: siScramble, siScramble + CDDP (15μM, 16 hours), siMAT2A (48 hours), and siMAT2A + CDDP in OVCAR3 cells.

**(H)** Overlap between DMRs identified from EPIC and differentially expressed genes from RNA-seq[30] after treatment with CDDP classified into four groups: genes exhibiting hypermethylation concurrent with transcriptional downregulation (Hyper+DOWN), hypomethylation with upregulation (Hypo+UP), Hyper+UP and Hypo+DOWN. OVCAR3 RNA seq and CpG probe overlap.

**Supplementary Figure S3. Inhibiting MAT2A increases chemotherapy sensitivity by enhancing DNA damage and reducing DNA repair in OVCAR5**

**(A-B)** Representative apoptosis assay plots (A) and quantification (B) of %apoptotic cells in OVCAR5 cells treated with siMAT2A-1 (24 hours), siMAT2A-3 (24 hours), platinum (cisplatin, CDDP, 12μM,16 hours), or combination.

**(C-D)** Representative comet assay images (C) and quantification (D) of tail moment as a measure of DNA damage in OVCAR5 cells treated with siMAT2A-1 (24 hours), siMAT2A-3 (24 hours), CDDP (12μM,16 hours), or combination. Box plots summarize the distribution of tail moment values for each treatment group; the center line indicates the median, the box represents the interquartile range, and the whiskers indicate the minimum and maximum values.

**(E-G)** Representative immunofluorescence images (E) and quantification of S9.6 (F) and γH2AX (G) in OVCAR5 wild-type and shMAT2A cells with or without CDDP treatment. Each dot represents the fluorescence intensity of a single cell, and the horizontal-colored line indicates the mean.

**(H)** Representative western blot of FANCD2 in OVCAR5 wild-type and shMAT2A cells with or without platinum treatment, with densitometric quantification normalized to the corresponding tubulin.

**(I)** Representative western blot of γH2AX in OVCAR3 treated with siMAT2A-1 (24 hours), siMAT2A-3 (24 hours), CDDP (12μM,16 hours), or combination, with densitometric quantification normalized to the corresponding H3.

**(J-K)** Representative scatter plots and quantification of HR-mCherry and NHEJ-GFP reporter activity in Kuramochi-DDR (J) and OVCAR5-DDR (K) cells with or without MAT2A shRNA knockdown. Each dot represents an independent biological replicate, and lines denote paired measurements obtained from the same biological replicate.

**(L-M)** Representative western blot of HR (L) and NHEJ (M) pathway proteins (phosphorylated and total) in OVCAR5 under indicated treatment conditions, with densitometric quantification relative to actin or tubulin from the corresponding sample.

**(N)** Representative western blot of NHEJ pathway proteins in OVCAR3 under indicated treatment conditions, with densitometric quantification relative to actin or tubulin from the corresponding sample.

**Supplementary Figure S4. SAM synthesis mediates DNA repair activation in response to platinum treatment through mTOR-S6K-FANCD axis in OVCAR5**

**(A)** Representative western blot of mTOR and p-S6K in OVCAR3 cells treated with platinum (cisplatin, CDDP; 15µM, 16 hours), and/or FIDAS-5 (11.8µM, 48 hours) or siMAT2A (48 hours).

**(B)** Representative western blot of mTOR and p-S6K in OVCAR5 cells treated with CDDP (12µM, 16 hours), and/or FIDAS-5 (10.9µM, 48 hours) or siMAT2A (24 hours).

**(C)** Co-immunoprecipitation (Co-IP) of FLAG-DEPDEC5-transfected OVCAR5 cells treated with siMAT2A with SAM concentrations (250 or 500 µM) added at lysate for overnight incubation, and densitometric quantification of BMT2 normalized to Flag-Depdc5.

**(D-E)** Western blot of MAT2A in OVCAR3 (D) and OVCAR5 (E) cells treated with CDDP (12µM, 16 hours), siMAT2A (48hours), or combination with or without SAM supplementation, with densitometric quantification normalized to tubulin.

**(F-J)** Western blot of mTOR and p70-S6K (F), FANCD2 (G), HR DNA repair proteins (H-I) and NHEJ DNA repair proteins (J) in OVCAR5 cells treated with CDDP (12µM, 16 hours), siMAT2A (48hours), or combination with or without SAM supplementation, with densitometric quantification normalized to actin.

**(K)** Representative comet assay images and quantification of tail moment in OVCAR5 cells treated with CDDP (12µM, 16 hours), siMAT2A (48hours), or combination with or without SAM supplementation.

**Supplementary Figure S5. Met- and MAT2A inhibitor prevented platinum-induced OCSC enrichment in OVCAR5**

**(A)** Representative immunohistochemical staining of HE (left), ALDH1A1 (middle) and SIRT1 (right) in xenograft tumors from vehicle- and CDDP-treated mice. ALDH1A1 and SIRT1 staining intensity was quantified using QuPath and is shown on the right. Scale bar = 50μm. Data are presented as mean ± SEM.

**(B-C)** Flow gating strategy and quantification of %ALDH+ cells (A) and representative brightfield images and quantification of spheroid (B) in OVCAR3 cells treated with platinum, Met(–), or Met(–) + platinum (cisplatin, CDDP; 12μM, 16 hours).

**(D-E)** Quantification of %ALDH+ cells (C) and representative brightfield images and quantification of spheroid (D) in OVCAR5 cells treated with CDDP, Met(–), or Met(–) + CDDP (12μM, 16 hours).

**(F)** Percentage of ALDH+ cells in OVCAR5 cells supplemented with CDDP (12μM,16 hours) or exogenous SAM (25-200μM, 40 hours).

**(G)** ssGSEA-derived composite scores (OCSC and general CSC hallmarks) and plotted against ΔMAT2A expression between matched recurrent and primary tumors.

**(H-I)** Percentage of ALDH+ cell (G), and representative images and quantification of spheroid formation (H) in OVCAR5 cells treated FIDAS-5 (10.9μM,48 hours) , CDDP (12μM,16 hours) or combination.

**Supplementary Figure S6. MAT2A siRNA prevented platinum-induced OCSC enrichment and SAM rescued the OCSC phenotype**

**(A-B)** Quantification of %ALDH+ cells (A) and representative images and quantification of spheroid formation (B) in OVCAR3 cells transfected with siScramble, siMAT2A-1, or siMAT2A-3 (48 hours) with or without platinum (cisplatin, CDDP; 15μM,16 hours)

**(C-D)** Quantification of %ALDH+ cells (C) and representative images and quantification of spheroid formation (D) in OVCAR5 treated with siScramble, siMAT2A-1, or siMAT2A (24 hours) with or without CDDP (12μM,16 hours)

**(E)** Quantification of %ALDH+ cells in Kuramochi treated with siScramble, siMAT2A-1, or siMAT2A-3 (48 hours) with or without CDDP (12μM,16 hours)

**(F)** Percentage of ALDH+ cells in OVCAR5 cells treated with CDDP (12μM,16 hours), siMAT2A (24 hours), or combination with or without 50 μM exogenous SAM (40 hours).

**(G)** Stemness marker protein expression by western blot with densitometric quantification to actin or tubulin in OVCAR5 cells treated with CDDP (12μM,16 hours), siMAT2A (24 hours), or combination with or without 50 μM exogenous SAM (40 hours).

**(H)** Representative images and quantification of spheroid formation in OVCAR5 cells treated with CDDP (12μM,16 hours), siMAT2A (24 hours), or combination with or without 50 μM exogenous SAM (40 hours).

(* $p<0.05$, ** $p<0.01$, *** $p<0.001$, **** $p<0.0001$; ns, not significant)

**Graphical Abstract: Mechanistic model of MAT2A-mediated platinum resistance in HGSC.** (Upper) Intact MAT2A activity supports DNA repair and SAM-dependent hypermethylation in response to platinum treatment, promoting CSC enrichment and cell survival. (Lower) MAT2A inhibition disrupts the mTOR-S6K-FANCD2 axis, causing R-loop accumulation and DNA repair failure. This, combined with the loss of platinum-induced hypermethylation, prevents stemness and restores platinum sensitivity to induce cell death.

#### Supplemental Figure S1

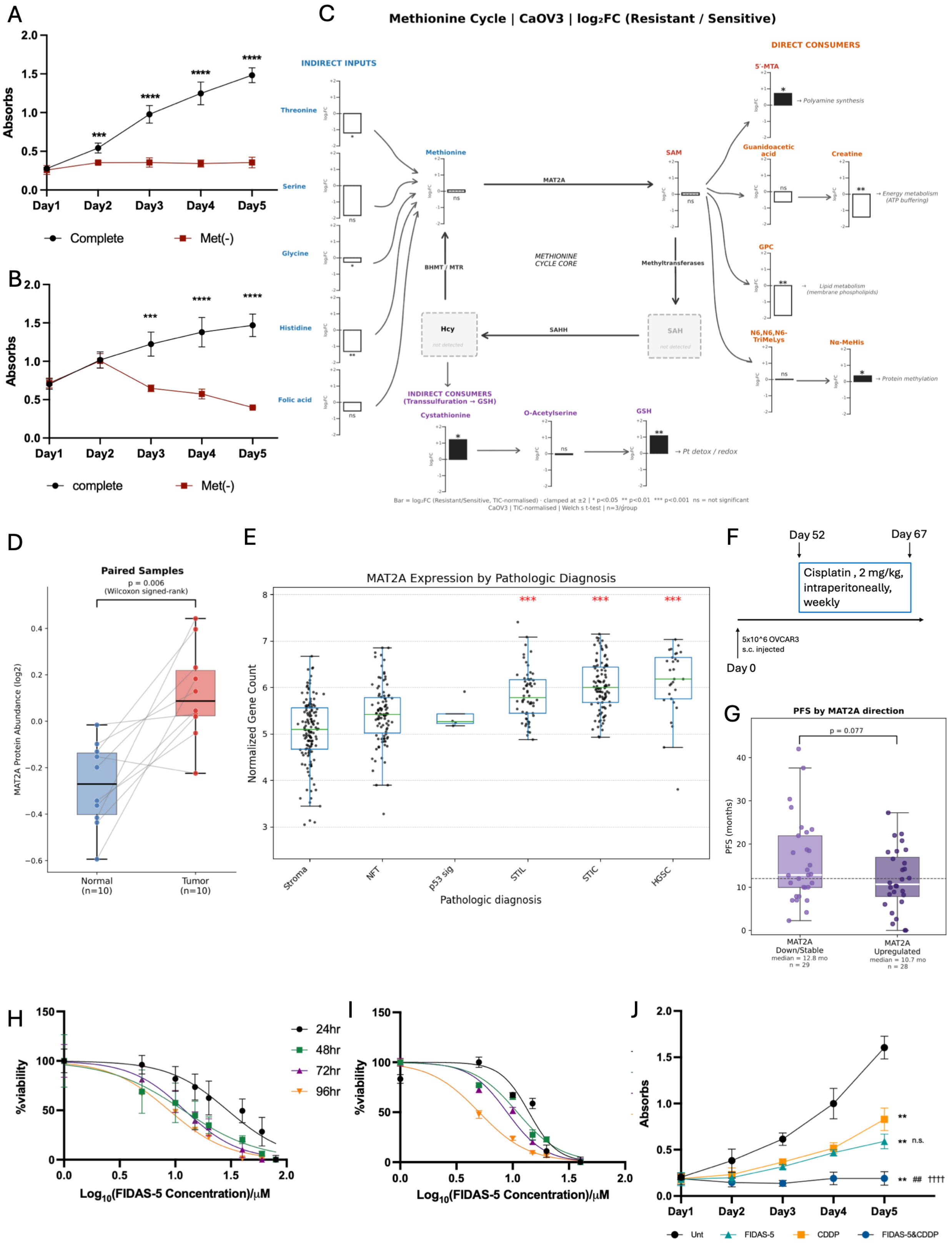

### Supplementary Figure S2

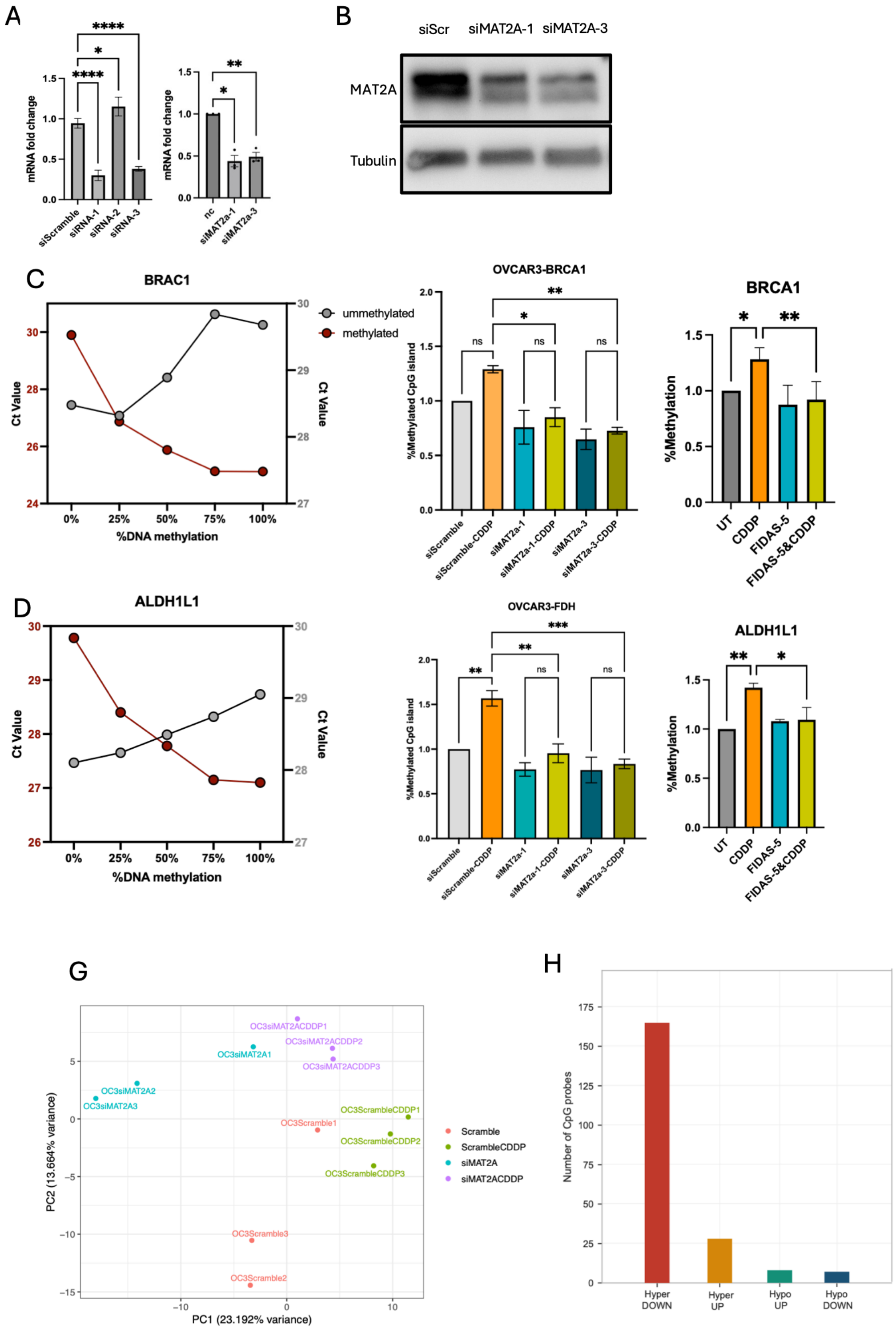

### Supplemental Figure S3

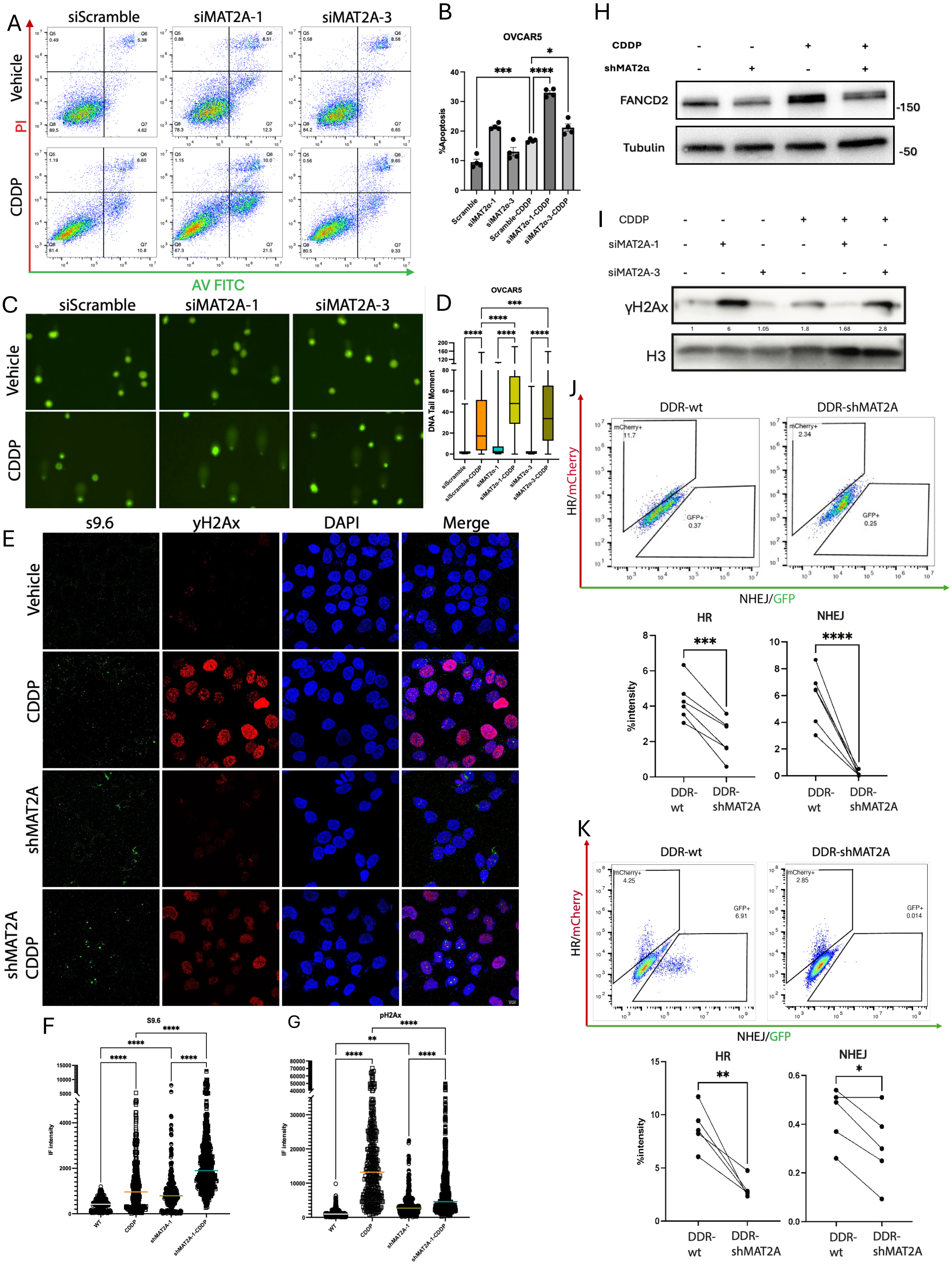

#### Supplementary Figure S3 Cont.

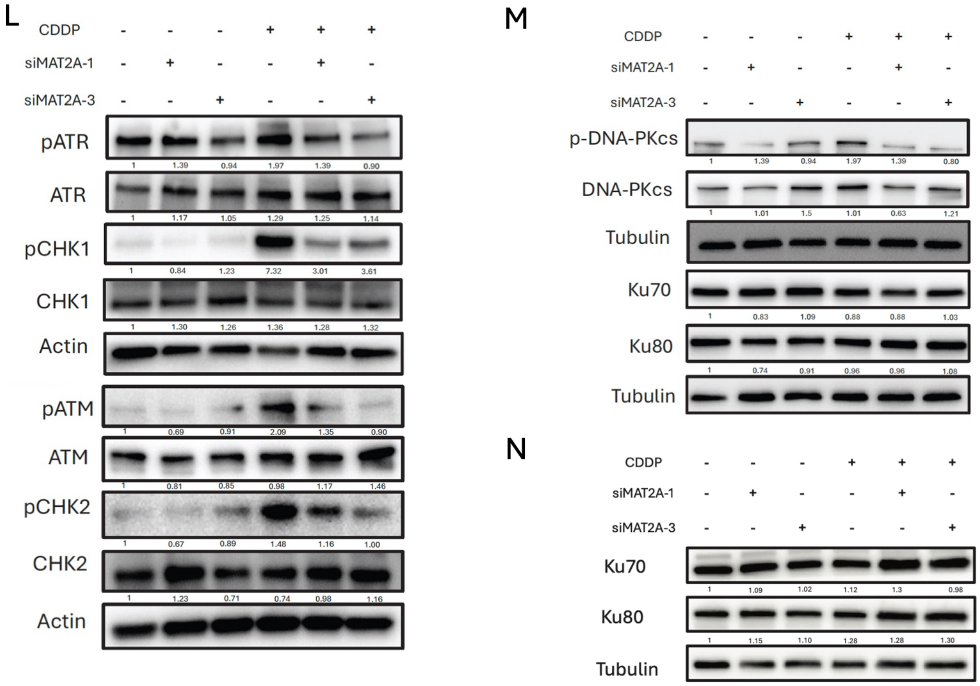

#### Supplemental Figure 4

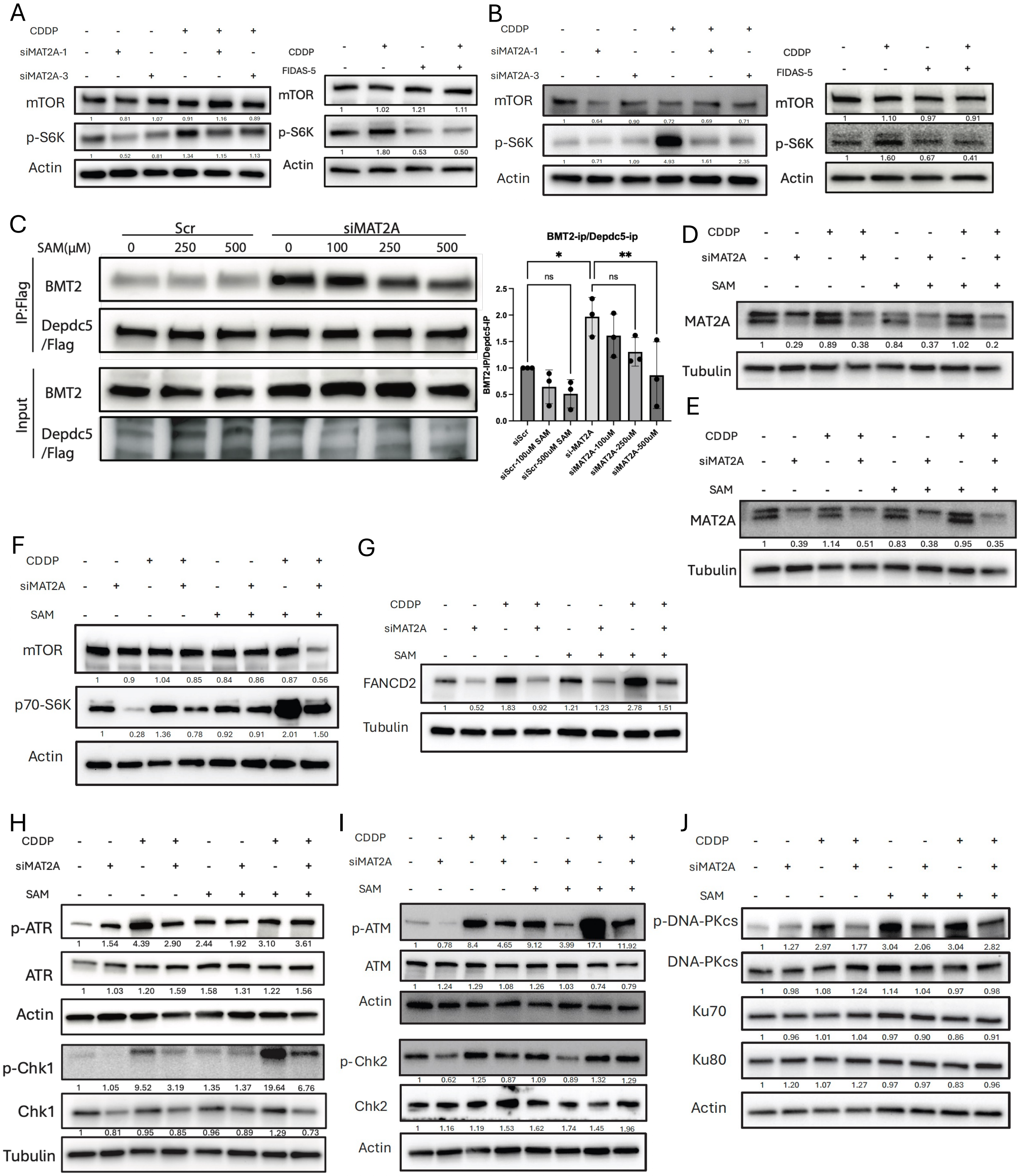

**Supplementary Figure S4. Cont.**

K

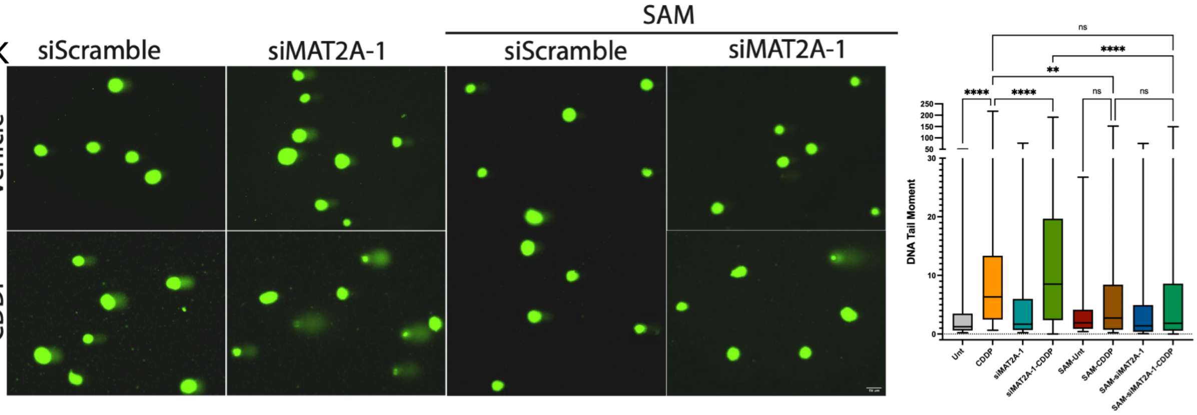

#### Supplemental Figure S5

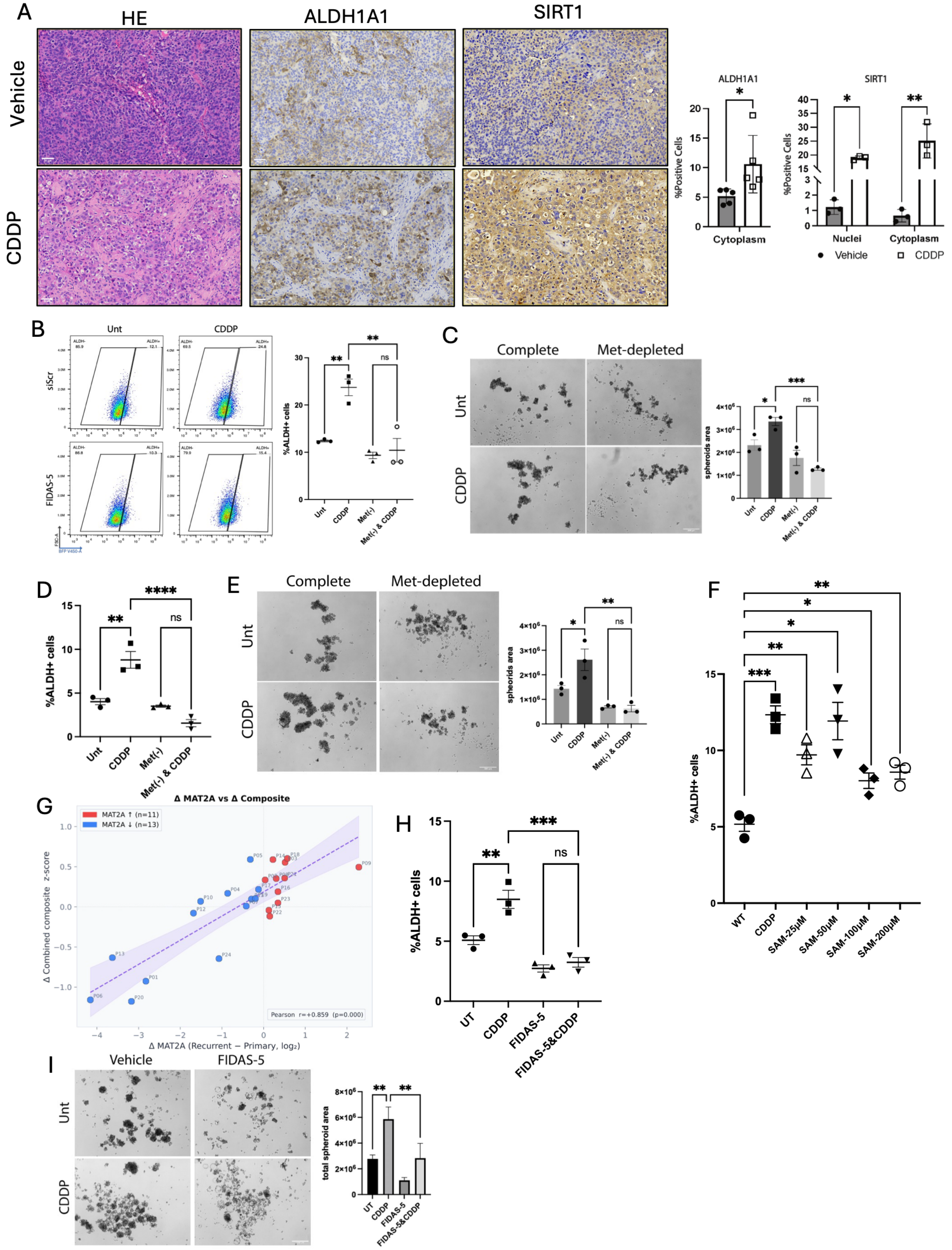

#### Supplemental Figure S6

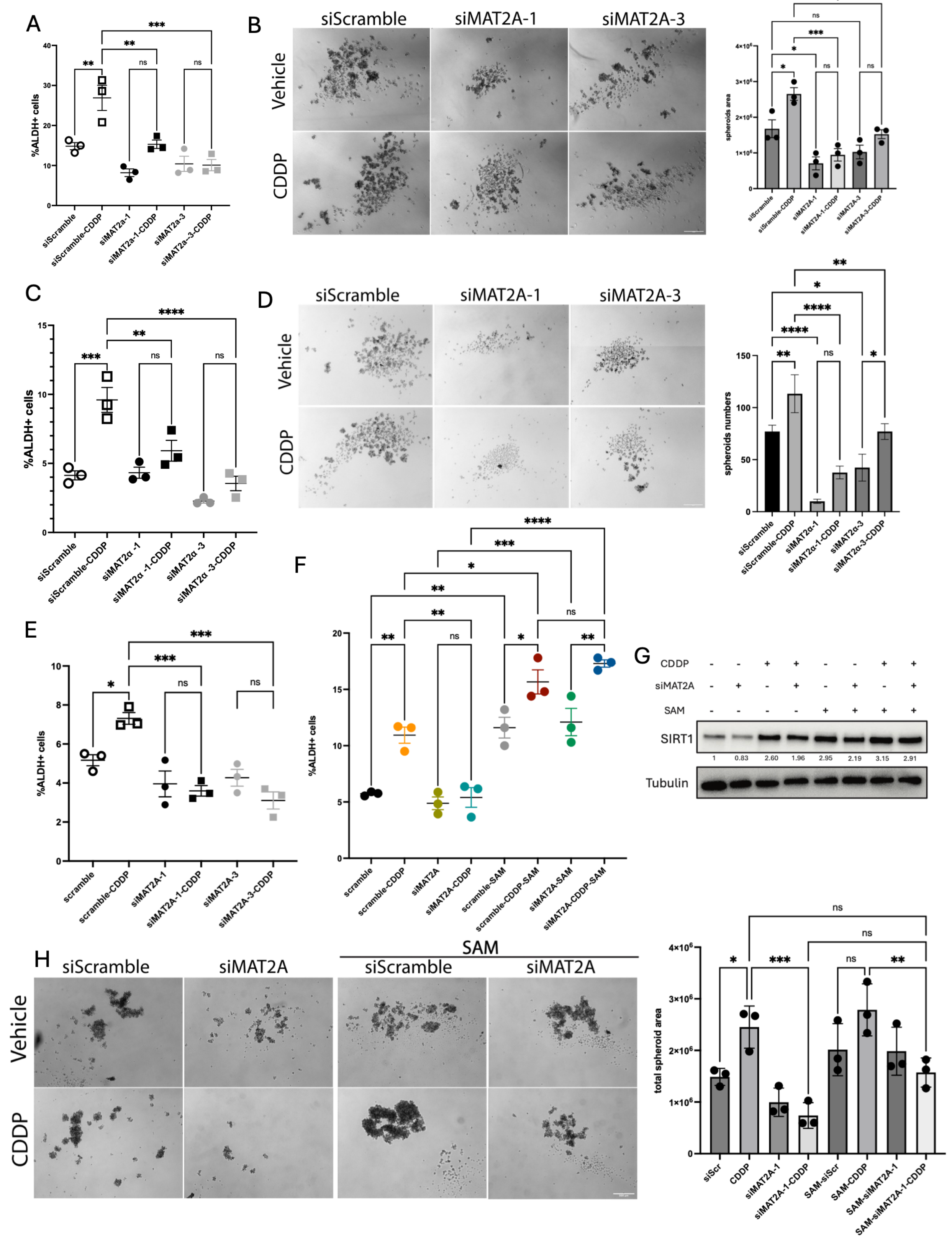

#### Graphical Abstract

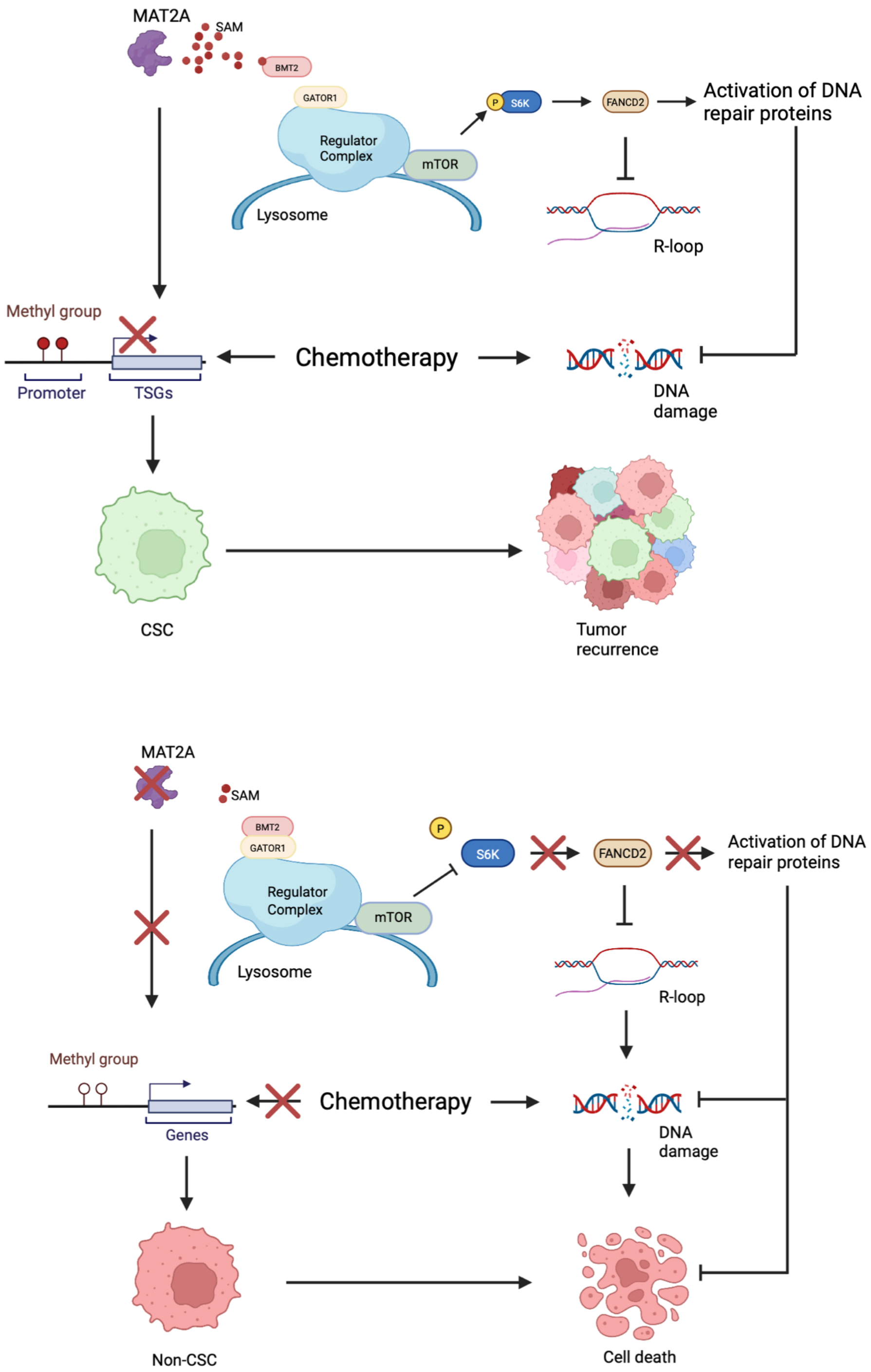
